# Identification of a multi-omics factor predictive of long COVID in the IMPACC study

**DOI:** 10.1101/2025.02.12.637926

**Authors:** Gisela Gabernet, Jessica Maciuch, Jeremy P. Gygi, John F. Moore, Annmarie Hoch, Caitlin Syphurs, Tianyi Chu, Naresh Doni Jayavelu, David B. Corry, Farrah Kheradmand, Lindsey R. Baden, Rafick-Pierre Sekaly, Grace A. McComsey, Elias K. Haddad, Charles B. Cairns, Nadine Rouphael, Ana Fernandez-Sesma, Viviana Simon, Jordan P. Metcalf, Nelson I Agudelo Higuita, Catherine L. Hough, William B. Messer, Mark M. Davis, Kari C. Nadeau, Bali Pulendran, Monica Kraft, Chris Bime, Elaine F. Reed, Joanna Schaenman, David J. Erle, Carolyn S. Calfee, Mark A. Atkinson, Scott C. Brackenridge, Esther Melamed, Albert C. Shaw, David A. Hafler, Al Ozonoff, Steven E. Bosinger, Walter Eckalbar, Holden T. Maecker, Seunghee Kim-Schulze, Hanno Steen, Florian Krammer, Kerstin Westendorf, IMPACC Network, Bjoern Peters, Slim Fourati, Matthew C. Altman, Ofer Levy, Kinga K. Smolen, Ruth R. Montgomery, Joann Diray-Arce, Steven H. Kleinstein, Leying Guan, Lauren I. R. Ehrlich

## Abstract

Following SARS-CoV-2 infection, ∼10-35% of COVID-19 patients experience long COVID (LC), in which often debilitating symptoms persist for at least three months. Elucidating the biologic underpinnings of LC could identify therapeutic opportunities. We utilized machine learning methods on biologic analytes and patient reported outcome surveys provided over 12 months after hospital discharge from >500 hospitalized COVID-19 patients in the IMPACC cohort to identify a multi-omics “recovery factor”. IMPACC participants who experienced LC had lower recovery factor scores compared to participants without LC. Biologic characterization revealed increased levels of plasma proteins associated with inflammation, elevated transcriptional signatures of heme metabolism, and decreased androgenic steroids in LC patients. The recovery factor was also associated with altered circulating immune cell frequencies. Notably, recovery factor scores were predictive of LC occurrence in patients as early as hospital admission, irrespective of acute disease severity. Thus, the recovery factor identifies patients at risk of LC early after SARS-CoV-2 infection and reveals LC biomarkers and potential treatment targets.

## INTRODUCTION

Long COVID (LC) has become a pressing public health concern, affecting an estimated 10-35% of surviving individuals infected with SARS-CoV-2, or 15-20 million individuals in the United States and over 60 million worldwide^1,2^. In July 2024, the National Academies of Sciences, Engineering, and Medicine (NASEM) released an updated definition of LC, characterizing it as a chronic condition arising after SARS-CoV-2 infection that persists for at least 3 months, irrespective of acute disease severity^2^. LC can encompass a wide range of physical and cognitive symptoms, and can lead to new or worsening neurological, psychiatric, cardiovascular, pulmonary, endocrine, and gastrointestinal conditions, among others^1–12^.

Several studies have identified demographic and clinical risk factors for LC^13–15^, including age^16–18^, female sex^16–21^, and longer hospital stays^18,22^. Higher viral loads^18,19^ and lower anti-SARS-CoV-2 antibody titers^18,23,24^ during the acute infection phase have also been associated with LC development. There are multiple, potentially overlapping hypotheses that explain the etiology of this condition, including persistent viral infection^25,26^, chronic inflammation^26–29^, latent herpesvirus reactivation^26,30,31^, immune dysregulation^29,32,33^, complement dysregulation^34^, and autoimmunity^35,36^. Despite these efforts, no consensus exists on the mechanisms of LC pathogenesis, and validation of the molecular findings across cohorts has been challenging. Additionally, most existing studies rely on measurements from a single or limited number of assays and are confined to restricted sampling time points during the acute or convalescent phases of the disease. Therefore, a multi-omics longitudinal study spanning both acute infection and convalescent disease phases could help elucidate the molecular mechanisms underlying LC.

The Immunophenotyping Assessment in a COVID-19 Cohort (IMPACC) study^37^ offers a unique opportunity to investigate the temporal dynamics of multi-omics immune profiles during the acute and convalescent COVID-19 infection phases in a clinically well-characterized cohort of hospitalized patients from across the United States. Data from the IMPACC study can be leveraged to identify molecular correlates of post-acute symptom development or resolution for one year after hospital discharge. This cohort has previously been studied to characterize multi-omics determinants of COVID-19 severity and mortality during the acute phase of disease^38–40^. Another study of the IMPACC cohort identified LC participants who experienced patient-reported outcome deficits up to 12 months after COVID-19 hospital discharge^18^. Clinical characteristics such as female sex, a higher respiratory SARS-CoV-2 viral burden, and lower antibody titers against the SARS-CoV-2 Spike protein during acute disease were associated with persistent deficits after hospital discharge. B cell lymphopenia and elevated fibroblast growth factor 21 (FGF21) during the acute disease phase were also characteristics of participants who developed LC^18^. However, longitudinal immune profiles of IMPACC participants experiencing LC during the convalescent phase of disease have not yet been compared to those of IMPACC participants who experienced minimal deficits during convalescence, an analysis that could reveal LC biomarkers and uncover biological processes underlying the disease.

In the current work, we applied supervised multi-omics integration methods to develop interpretable models that differentiate participants with LC from recovered individuals based on their longitudinal immunophenotyping profiles during the convalescent disease phase. We identified key biological programs and biomarkers driving LC classification. Our findings highlight persistent inflammation, dysregulation of heme metabolism, and altered androgenic steroid profiles as characterizing features of LC, independent of acute disease severity or SARS-CoV-2 vaccination status post hospital discharge. Notably, these molecular profiles were already detectable during the acute phase of disease, suggesting their potential value as early predictive biomarkers for identifying patients at risk of developing LC. Additionally, despite a general lack of consensus about the definition of LC or consistency in the timing of sampling across different studies, we validated dysregulation of the heme metabolism signature in an independent LC cohort. These findings provide valuable insights into the molecular underpinnings of LC and offer a foundation for future research aimed at improving diagnostics and developing targeted interventions.

## RESULTS

### Longitudinal multi-omics profiling of long COVID

The IMPACC study included 1,164 participants admitted to 20 US hospitals for COVID-19 infection between May 2020 and March 2021^37^. Clinical data collection and immunophenotyping were performed longitudinally during the acute disease phase within 72h of hospital admission and 4, 7, 14, 21 and 28 days after hospital admission (Visits 1-6, respectively)^2^. Surviving participants were contacted 3, 6, 9, and 12 months after hospital discharge (Visits 7-10, respectively) to complete patient-reported outcome and symptom surveys during the convalescent phase, and to provide biosamples for immunophenotyping assays. Of the 702 participants who could be reached by the study team after discharge, 513 were included in the IMPACC Convalescent cohort. These participants were selected because they completed at least one survey and provided at least one biosample during the convalescent period (Figure 1A and Table S1)^18^. IMPACC core labs performed immunophenotyping both in the acute and convalescent phases, including measurements of inflammatory mediators in blood serum via Olink (SO), global blood plasma metabolomics (PMG), global and targeted blood plasma proteomics (PPG and PPT), peripheral blood mononuclear cell (PBMC) transcriptomics (PGX), whole blood cell frequencies measured by mass cytometry by time of flight (CyTOF), and CyTOF mean marker signal intensity measurements (BCT).

**Figure 1.**
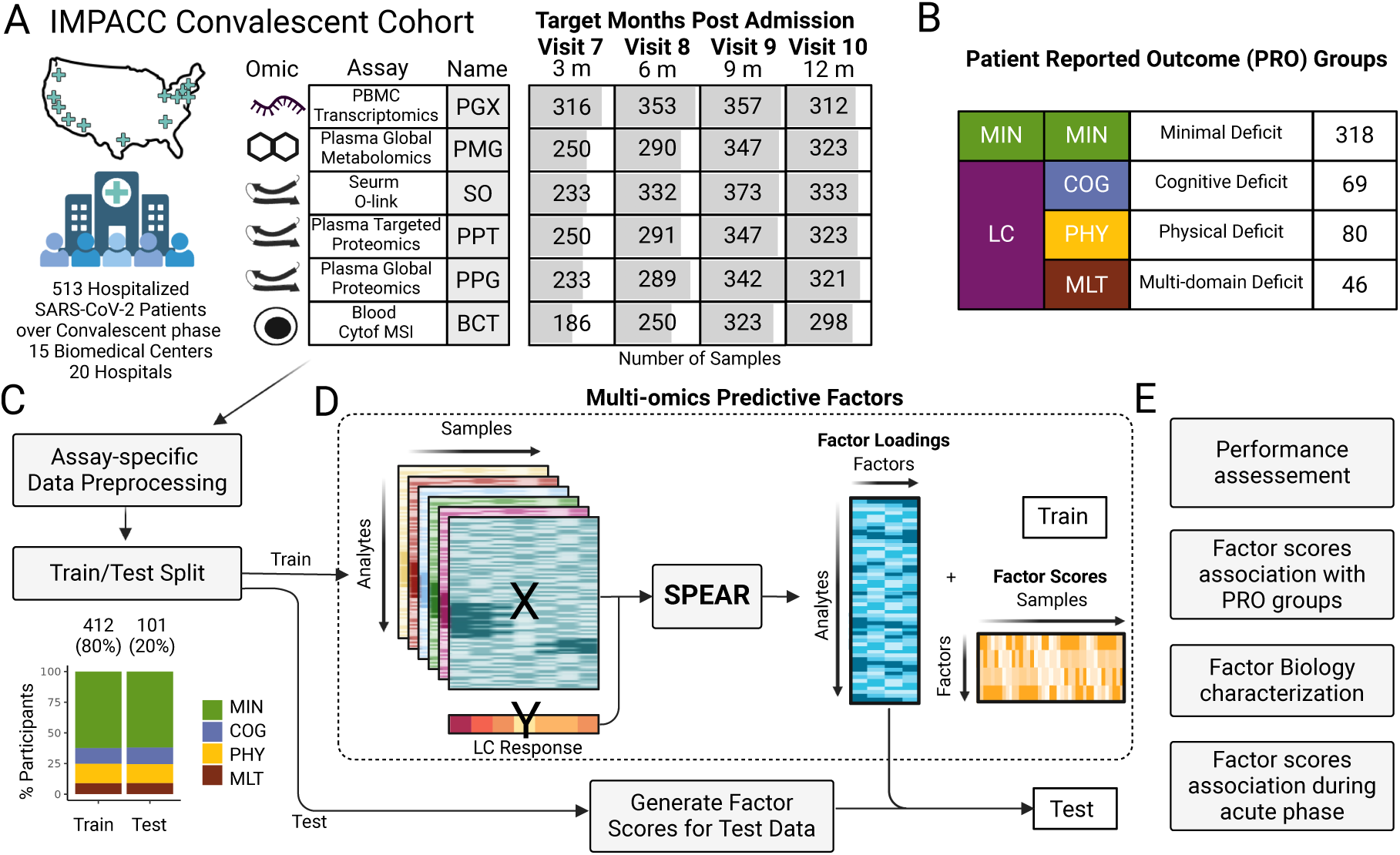
Multi-omics data overview and generation of a predictive LC factor. **(A)** Number of samples used in the multi-omics data integration strategy by assay (rows) and scheduled time of collection (columns). Shading indicates the frequency of samples with data availability at the indicated visit. **(B)** Patient classification in Patient Reported Outcome (PRO) clusters according to the PRO measure survey scores^18^. **(C)** Individual assay data were preprocessed and split into Train and Test cohorts by participant in an 80/20 split, maintaining the proportion of PRO cluster participants in each partition. **(D)** Preprocessed assay data and LC response outcomes for the Train cohort were used to identify multi-omics predictive factors with SPEAR. Factor scores were then calculated for the Test cohort. **(E)** The performance of the multi-omics predictive factors to classify patients into presence and absence of LC was assessed on the Train cohort via cross-validation and then validated on the Test cohort. The predictive factor scores were confirmed to be associated with LC after correcting for possible confounding variables. In depth analysis of enriched biological pathways and significant analytes relevant for the prediction was performed. Factor scores were computed for the acute infection immune profiles, and association analysis with LC at these early time points was performed. See also Figure S1.

LC status was defined in this cohort according to the participant’s response to post discharge surveys that captured symptoms and Patient-Reported Outcome (PRO) measures that evaluated general health and deficits in specific domains. Participants who responded to at least one set of post discharge surveys were assigned to PRO clusters according to latent class modeling and clustering using standardized scores of the PRO measures, as previously reported^18^ (details on the PRO measures used can be found in the Supplementary Methods). PRO clusters were classified as participant clusters with no or minimal deficits (MIN), or with deficits attributed to LC in several domains: physical predominant (PHY), mental/cognitive predominant (COG), and multi/pan domain (MLT)^18^ (Figure 1B).

In this study, we utilized multi-omics immunophenotyping profiles to develop interpretable models for predicting LC and exploring the underlying molecular mechanisms. To assess the model performance, we split the Convalescent cohort into an 80% Train and 20% Test cohort, maintaining the proportions of participants in each PRO cluster (Figure 1C), with no noticeable imbalance in other clinical characteristics or biosample availability between the cohorts (Figure S1). We then used Signature-based multiPle-omics intEgration via lAtent factoRs (SPEAR)^41^, a supervised Bayesian factor model for the identification of multi-omics features, to integrate the high dimensional data and construct multi-omics predictive factors from immune profiles obtained during the convalescent phase in the Train cohort. We assessed their predictive performance by repeated cross-validation on the Train cohort and validated the selected model performance on the Test cohort (Figure 1D and 1E). To identify the immune programs captured in the predictive factors, we conducted in depth analyses of enriched biological pathways and analytes identified as highly relevant for the prediction performance by the model and performed associations with assay data not included in model training, such as blood CyTOF cell frequencies.

### Multi-omics factors are predictive of long COVID

We focused on predicting LC in the Convalescent cohort from the multi-omics immune profiling data collected during the convalescent phase. Since the binary LC labels (presence or absence of LC) per participant could omit valuable information captured by the numeric PRO measures at each participant visit, we constructed separate SPEAR models to generate supervised factors including PRO measure scores (SPEAR Physical, SPEAR Cognitive, SPEAR Mental, SPEAR Impact, SPEAR Dyspnea) or the LC binary labels (SPEAR LC) as response variables (Figure S2A). We evaluated the predictive performance of the SPEAR multi-omics factors trained on the individual PRO measure scores to reconstruct the score values of unseen data in the training cohort by cross-validation using a lasso regression model (Figure S2B). The SPEAR Physical model was the best performing, with lowest prediction error, and was selected for further evaluation. We then utilized the same procedure to assess the ability of the SPEAR LC factors and the best performing SPEAR model trained on individual PRO measure scores (SPEAR Physical) to predict binary LC status. The lasso model trained on the SPEAR Physical multi-omics factor achieved the highest predictive performance as evaluated with the area under the receiver-operating characteristic curve (AUROC) (Figure 2A). Additionally, we compared the predictive performance of the SPEAR multi-omics supervised factors to the performance of equivalent unsupervised multi-omics factors, obtained with the Multi-Omics Factor Analysis (MOFA)^42^ framework, that do not consider a response variable during the factor construction step. Both lasso classifiers trained on the SPEAR Physical and SPEAR LC predictive factors outperformed the classifier using MOFA unsupervised factors on the Train cohort (Figure 2A, Figure S2B). The SPEAR Physical model achieved an AUROC of 0.69 for predicting LC presence or absence in the Test cohort (Figure 2B). The SPEAR Physical Factor, learned by the SPEAR physical model, was significantly associated with LC in the Test cohort after correcting for sex and age (p=0.00098), two variables which have been previously associated with LC in our cohort^18^. The SPEAR Physical Factor scores were significantly higher for participants in the MIN group compared to the LC group, so we termed this factor as the “recovery factor” (Figure 2C, Figure S4A). Recovery factor scores were significantly associated with PRO clusters (p=0.0009); however, they showed a differential ability to identify individual LC deficit domains, with significant differences between MIN vs COG and MIN vs MLT PRO clusters, but not MIN vs PHY PRO clusters (Figure 2D, Figure S4B). Sparse lasso regression models to reconstruct recovery factor scores utilizing all analytes included in the model or analytes from individual omics assays showed that the model including all assays was best at reconstructing the factor scores, indicating that the recovery factor scores captured contributions from multiple omics layers (Figure S2C). Taken together, the recovery factor is a multi-omics model comprised of biologic analyte levels during the convalescent phase of COVID-19 that is able to distinguish MIN from LC over a 12 month period post hospital discharge in the IMPACC cohort.

**Figure 2.**
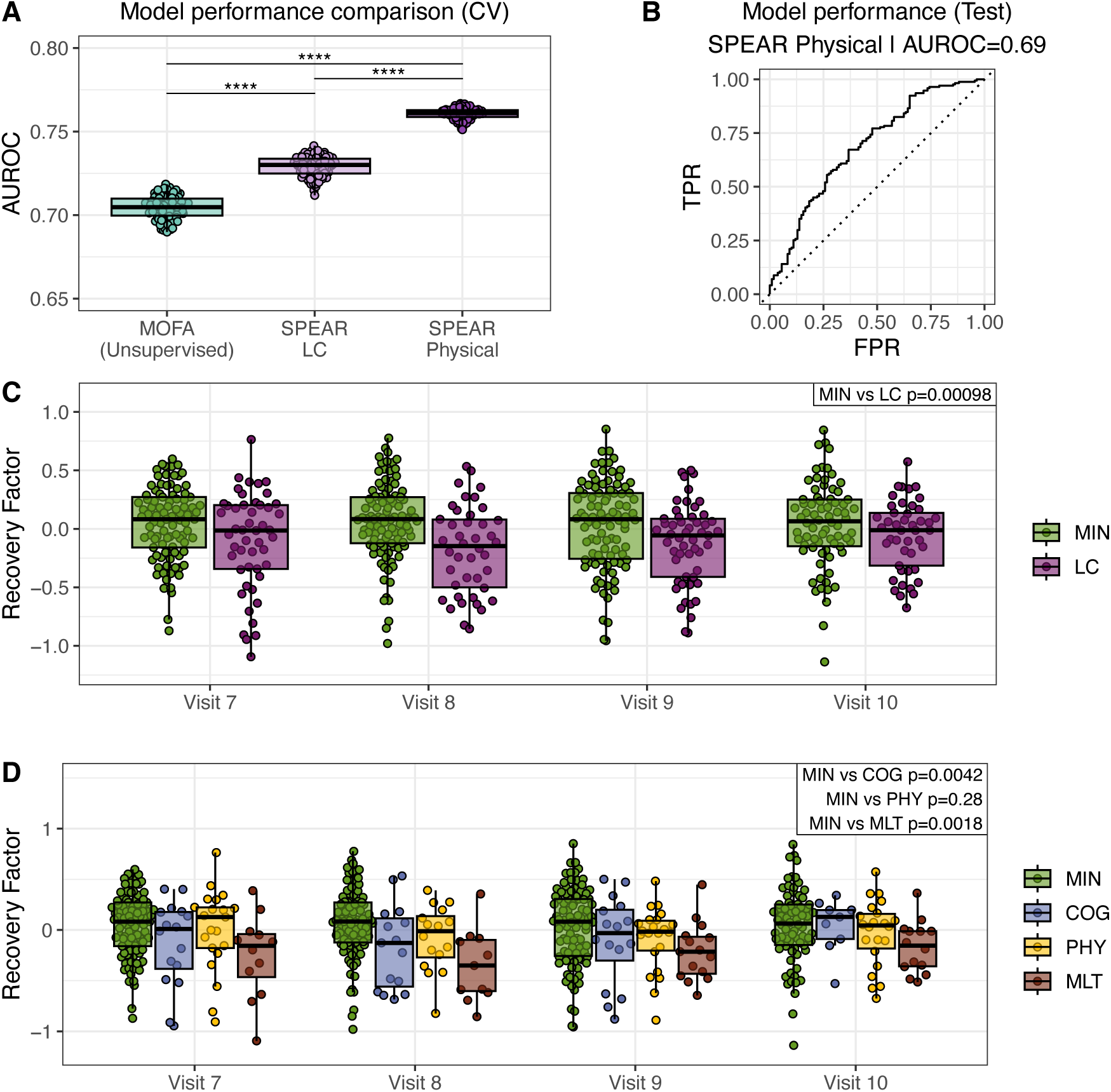
Identification of a convalescent multi-omics recovery factor that discriminates long COVID. **(A)** Predictive performance of a lasso model trained on the MOFA and SPEAR factors to discriminate LC vs MIN at the event level. The mean AUROC of a 10-fold cross-validation on the Train Cohort, for 100 bootstrapped model training repetitions are shown. Significance was calculated by standard normal approximation of bootstrapped differences between models (t-test, ****adj. p-value ≤ 0.0001) **(B)** Predictive performance of the SPEAR Physical model to discriminate LC vs MIN on the Test cohort. ROC curve of model (solid line), random classifier (dashed line), and AUROC value are shown. TPR: true positive rate, FPR: false positive rate. **(C)** Recovery factor scores for the Test cohort of the MIN and LC groups at 3 months (Visit 7), 6 months (Visit 8), 9 months (Visit 9) and 12 months (Visit 10) after hospital discharge. **(D)** Recovery factor scores of the individual PRO clusters by visit for the Test cohort. P-values in C and D show the significance of the recovery factor score association with MIN vs LC and pairwise PRO cluster combinations, respectively (see methods for association details). See also Figure S2, S3, and S4.

### Functional characterization of the recovery factor

To characterize the biologic processes underlying the recovery factor, we performed gene set enrichment analysis (GSEA) for each of the multi-omics assays based on the SPEAR model’s internal ranking of the relative importance of each feature for predicting the PRO Physical score. Surveyed pathways included gene sets from the Molecular Signatures Database (MSigDB) Hallmark^43^ and KEGG^44^ resources as well as metabolite sets from the Subpathway resource^45–47^. Joint p-values were computed to assess analyte set enrichment across the multi-omics assays. The Hallmark Heme Metabolism transcriptomic pathway was negatively associated with the recovery factor, indicating upregulation in LC participants, whereas the androgenic steroids metabolite set was positively associated with the recovery factor, indicating downregulation in LC participants (Figure 3A). Evaluated individually, several leading edge analytes in the Hallmark Heme Metabolism gene set and androgenic steroids Subpathway metabolite set showed significant associations with LC status (Figure S5 A and B).

**Figure 3.**
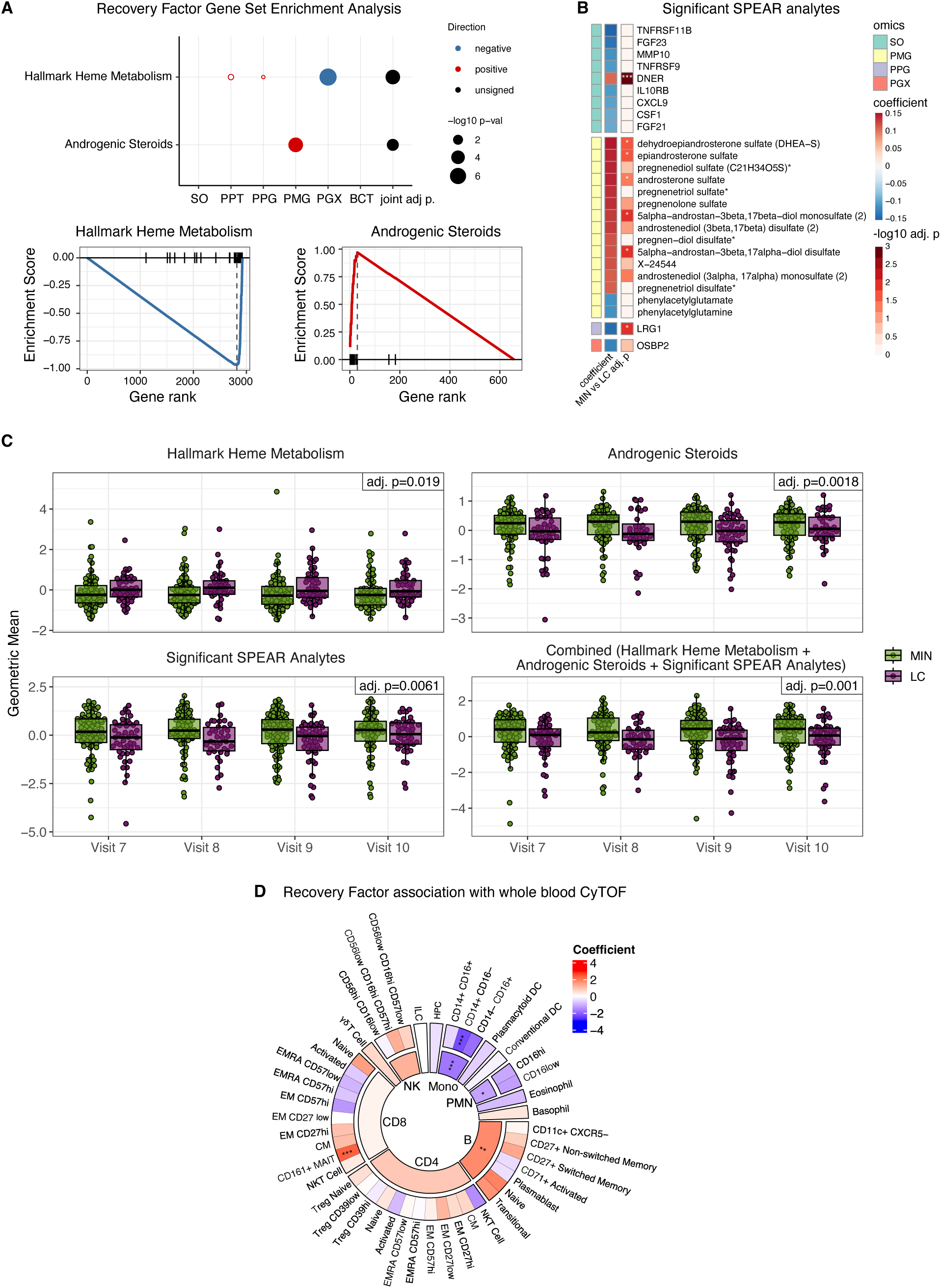
Heme metabolism and androgenic steroid pathways, inflammation-associated serum factors, and altered immune cell composition are associated with the recovery factor during convalescence. **(A)** GSEA identifies heme metabolism and androgenic steroid pathways as significantly associated with the recovery factor, with significance shown per assay, as well as across assays (joint adj. p < 0.05). **(B)** 26 significant analytes (SPEAR Bayesian posterior selection probability ≥ 0.95) in the recovery factor across different assays (left) were identified using SPEAR factor loadings (middle; coefficient in the factor), and each was tested for association in the test cohort with MIN vs LC groups (right; adj. intercept p-value). **(C)** Geometric means of analytes from the significantly enriched gene and metabolite sets and/or significant SPEAR analytes are shown per sample at each convalescent visit in the test cohort. The p-values indicate significance of the association with MIN vs LC. **(D)** Association in the full cohort of whole blood cell counts determined by CyTOF with the recovery factor for parent and child immune cell types. NK: Natural Killer cells, Mono: Monocytes, PMN: polymorphonuclear neutrophils, B: B lymphocytes, CD4: CD4+ T lymphocytes, CD8: CD8+ T lymphocytes. For a full list of the child populations see Table S3. (* adj. p-value < 0.05, ** adj. p-value < 0.01, *** adj. p-value < 0.001). See also Figure S5.

SPEAR performs internal significance testing to determine the importance of each analyte in predicting the response variable. The SPEAR Physical model identified 26 analytes across four assays that were significant in the recovery factor (SPEAR Bayesian posterior selection probability ≥ 0.95), and we performed individual associations of these features with LC status in the test cohort, adjusting for age and sex (Figure 3B, Figure S5). Nine of these 26 analytes were from the serum Olink assay. Of these, DNER (Delta And Notch-Like Epidermal Growth Factor-Related Receptor), a non-canonical Notch ligand that has been implicated in promoting tumor growth and metastasis and in supporting wound healing^48,49^ was significantly reduced in LC participants, consistent with a prior study of plasma proteomics in LC subjects^28^. The remaining serum Olink analytes were negatively associated with the recovery factor. In particular, they included proteins and cytokines associated with chronic inflammatory conditions^50–55^, particularly endothelial/vascular inflammation (FGF23, FGF21, CXCL9, TNFRSF11B and TNFRSF9 (CD137)), as well as inflammation-associated myeloid regulators^56–58^ (MMP10 and CSF1). Elevated levels of IL10RB have been previously associated with worse outcomes in acute COVID-19 infection^59^, consistent with elevation under inflammatory conditions. LRG1, a protein elevated in LC participants, is induced by IL-6 and other inflammatory cytokines and has been implicated in angiopathic activity^60–62^. Phenylacetylglutamate and phenylacetylglutamine are gut microbiota-derived metabolites associated with vascular inflammation and thrombosis^63^. Finally, the OSBP2 (ORP4) transcript, which encodes an oxysterol binding protein^64^, was a leading edge gene in the Hallmark Heme Metabolism gene set that was elevated in LC participants.

Several metabolites from the androgenic steroids pathway were represented in the 26 significant analytes and were positively associated with the recovery factor, indicating higher levels correlate with better physical function. When we tested these metabolites for their individual association with LC status, five (DHEA-S, epiandrosterone sulfate, androsterone sulfate, 5alpha-androstan-3beta,17beta-diol monosulfate (2), 5alpha-androstan-3beta,17alpha-diol disulfate) were significantly lower in LC participants, adjusting for age and sex (Figure 3B). Androgens can suppress inflammation^65^, suggesting that the higher level of androgenic steroids in participants of the MIN group could reflect better control of chronic inflammation. These findings are consistent with prior reports showing lower levels of sex hormones in LC^31^. Five metabolites related to pregnenolone were also represented in the significant SPEAR analytes (Figure 3B). Pregnenolone is synthesized from cholesterol as the first step of the steroid hormone biosynthesis pathway and is known to have potent effects as an inhibitor of inflammation^66^ and as a neurosteroid^67^. Altogether, these findings are consistent with a prominent role for persistent inflammation in LC with dysregulation of key analytes that may contribute to symptoms in LC, including elements that drive angiopathy, reduce wound healing, and alter heme metabolism.

The feature sets from heme metabolism and androgenic steroids identified by GSEA analysis combined with the significant SPEAR analytes represent 73 unique features that potentially condense the predictive power of the recovery factor into a smaller feature set. To test this hypothesis, we calculated the geometric mean of the 43 leading edge heme metabolism and 12 androgenic steroid features, as well as the 26 significant SPEAR analytes. All three geometric mean scores were independently significantly associated with LC in the test cohort (Figure 3C). Furthermore, the combined score that includes analytes from all three feature sets discriminates MIN and LC participants with even greater significance (Figure 3C). Thus, while the recovery factor is comprised of weighted contributions from 6,807 features, we have identified a smaller set of 73 unique features that discriminates participants according to LC status in the convalescent period.

Consistent with our finding, the Hallmark Heme Metabolism pathway was previously reported by Hanson et al.^29^ as an enriched pathway in participants with persisting symptoms 1-3 months after acute SARS-CoV-2 infection compared to participants without persisting symptoms. This cohort comprised 102 participants, including non-hospitalized and hospitalized individuals^29^. To determine whether the same heme metabolism-related genes were dysregulated in LC participants in the IMPACC and Hanson et al. cohorts, we used the leading edge genes from the significant Hallmark Heme Metabolism pathway in our GSEA analysis (Figure S5A) and calculated the geometric mean gene expression in PBMCs from the Hanson et al. cohort^29^. We found that our heme metabolism leading edge genes significantly differentiated participants with persistent vs. resolved symptoms after COVID-19 infection at multiple time points in the independent cohort (Figure S5C), validating the reproducibility of the gene expression datasets and underscoring the importance of this subset of heme metabolism genes.

Prior studies have identified altered leukocyte frequencies as a feature of LC^18,26,29,31,33^. To determine whether similar cellular changes were associated with the recovery factor, we analyzed whole blood CyTOF cell frequencies for 15 parent and 46 child immune cell types in our cohort during convalescence (Figure 3D). We found several cell subsets that were significantly associated with the recovery factor. B cells and CD161+ MAIT cells were positively associated with the recovery factor. In contrast, polymorphonuclear leukocytes (PMN) and monocytes, specifically the CD14^+^CD16^−^ classical monocyte subset, were negatively associated with the recovery factor. Together, these findings suggest that a persistent elevation in monocytes and neutrophils, along with a deficit in B cells, is associated with prolonged inflammation during LC. These findings are consistent with a previous report that monocytes are elevated in males with LC^31^. The decrease in MAIT cells with LC could be another effect of sustained inflammation as lower levels of circulating MAIT cells have been associated with chronic HIV^68^ and hepatitis C^69^ viral infections.

### The recovery factor is associated with clinical characteristics and multiple patient reported outcomes in the convalescent period

We next evaluated whether the recovery factor was associated with clinical features and additional clinical outcomes. We tested the association of recovery factor scores with clinical features at hospital admission (i.e., Visit 1), including demographics, comorbidities, complications, and baseline lab measurements (Figure 4A). Several demographic and clinical measures were significantly associated with recovery factor scores, including age, sex, length of hospital stay and the Sequential Organ Failure Assessment (SOFA) score. Notably, anemia as a complication was negatively associated with the recovery factor, whereas hemoglobin and hematocrit baseline measurements showed a significant positive association (Figure 4A). We additionally conducted association testing with Patient Reported Outcome (PRO) measures from surveys conducted at the same visit at which the recovery score was assessed in participants across the convalescent period, correcting for age and sex. The recovery factor score was significantly associated in the test cohort with the PROMIS Physical score, on which the model was trained (Figure 4B), and the EQ-5D-5L score, both of which contained similarly worded questions assessing physical function (Figure 4B). Interestingly, the recovery factor was also correlated with PROMIS Mental and PROMIS Psychosocial Impact scores, although these associations did not remain significant after p-value correction (Figure 4B). We also tested whether recovery factor scores associated with whether participants reported clinical symptoms in the 7 days prior to each visit but found no significance with any symptom group (Figure 4C).

**Figure 4.**
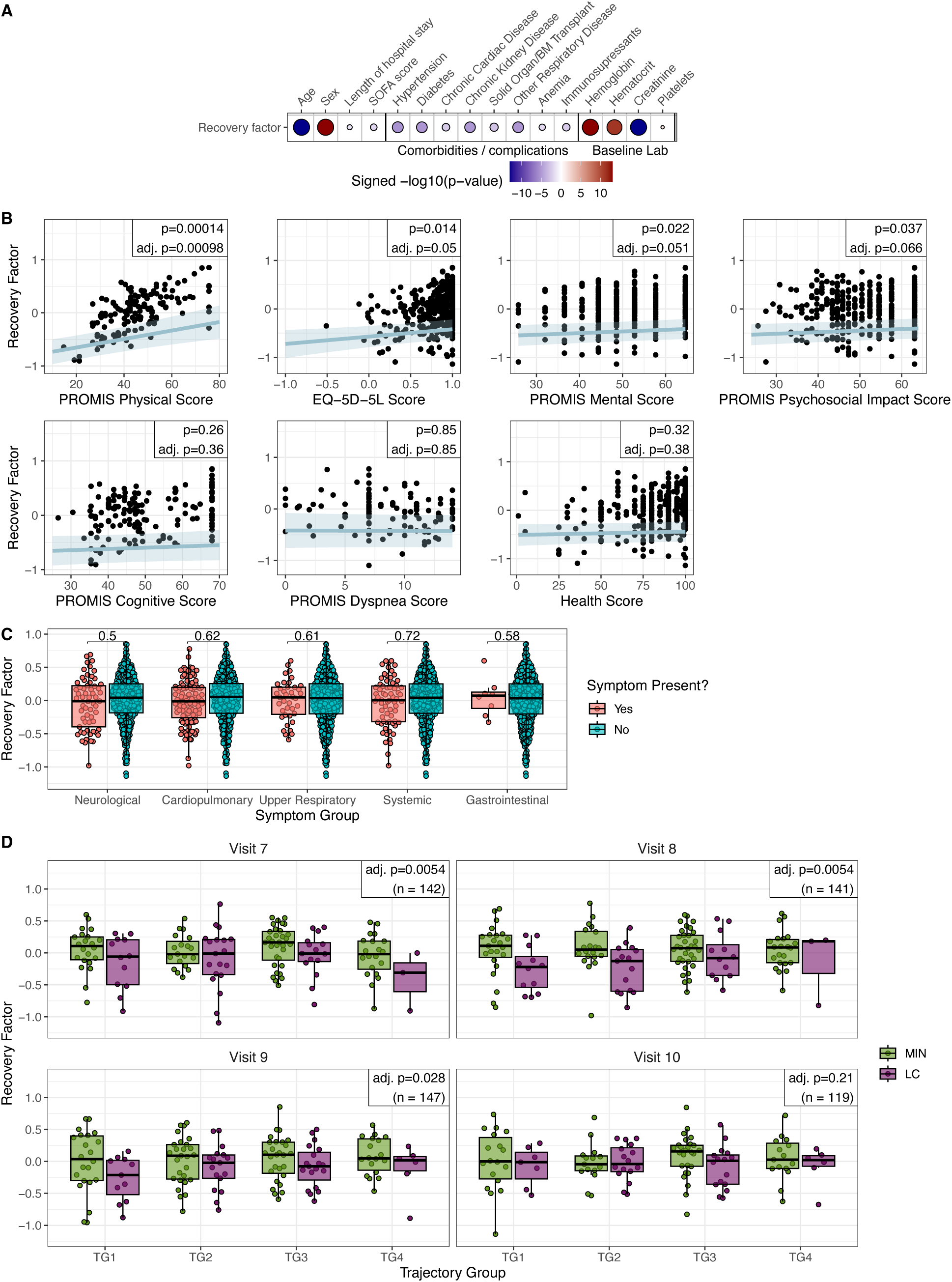
Associations of clinical measurements with recovery factor scores. **(A)** Association of recovery factor sores with clinical features (demographics, comorbidities, complications and baseline lab measurements). Dot plot shows the signed adjusted p-values indicating the clinical feature term significance from a linear mixed-effect model with enrollment site and participant as random effects to explain the convalescent phase recovery factor scores. Sex and discretized age were further adjusted as fixed effects for clinical features other than sex and age. Only significant associations (adj. p-value <0.05) are shown. **(B)** Associations of recovery factor scores with individual PRO survey scores (PROMIS scale scores, EQ-5D-5L and health score) in the test cohort. Raw and adjusted p-values indicated the PRO score term significance in linear mixed effect models. **(C)** Associations of recovery factor scores with each indicated symptom group in the test cohort. Numbers are the uncorrected significance (p-values) of the symptom group term in linear mixed effect models. **(D)** Recovery factor scores per participant in the test cohort, separated into MIN and LC groups by acute phase trajectory groups, stratified by visit. P-values for panels B-D show the endpoint term of a linear mixed effect model with sex, discretized admit age, and trajectory group as fixed effects and enrollment site as random effect. No individual MIN vs. LC comparisons were significant after p-value correction. (* adj. p-value < 0.05, ** adj. adj. p-value < 0.01, *** adj. p-value < 0.001)

There is a general lack of consensus about whether LC is associated with the severity of acute disease. A previous analysis of clinical features from the IMPACC cohort showed no association between the severity of acute infection, as assessed by clinical trajectory groups, and LC development^6,18^. However, other studies have found an association^6^. Thus, we sought to determine whether acute disease severity contributed to the association between recovery factor scores and LC status in our cohort. Clinical severity in the IMPACC cohort during the acute phase was defined by unsupervised clustering of respiratory ordinal score over time, taking discharge status and limitations into account, with trajectory group 1 (TG1) representing the mildest disease course and TG4 representing the most severe disease among participants who survived for at least 28 days post hospitalization^70^. After correcting for acute phase trajectory group assignment, recovery factor scores remain significantly associated with LC at the first three convalescent time points (Figure 4D), indicating that acute clinical severity does not contribute to the association between participant recovery factor scores in the convalescent phase of disease and LC status.

### Sex impacts recovery factor scores

The incidence of LC is higher in females than males, despite a higher percentage of males with severe COVID-19 acute disease courses^21,71^. In the IMPACC Convalescent cohort, nearly half of the female participants presented with long term deficits compared to only ∼30% of male participants (Fig. S6A). Assignment to clinical subtypes of LC was not influenced by sex, with similar proportions and numbers of male and female LC participants assigned to COG, PHY and MLT PRO clusters (Figure S6A). However, consistent with the known influence of sex on LC status, sex was a statistically significant covariate in the association of the recovery factor with LC status from Figure 2C (p=3.6e-7) and with PRO clusters from Figure 2D (adj. p<0.001 in all pairwise comparisons). Thus, we tested if the recovery factor was able to discriminate LC in both males and females by repeating our associations with LC status in the Test cohort separated by sex. Recovery factor scores were significantly associated with the binary assignment to LC vs. MIN groups in females but not in males after p-value adjustment (Figure S6B), although the trend of lower scores in LC participants persisted in males. When considering individual PRO groups, recovery factor scores discriminated between MIN vs. COG and MIN vs. MLT groups for females and MIN vs. MLT PRO groups for males (Figure S6C). Given that the incidence of LC is lower in males, it is notable that recovery factor scores were generally higher in males versus females regardless of LC status.

We next investigated whether the gene and metabolic pathways or top analytes associated with the recovery factor (Figure 3A-B) were differentially represented between sexes, where sex was also a statistically significant covariate (Figure 3C, adj. p<0.001). The geometric mean score of the heme metabolism pathway approached significance but was no longer significantly associated with LC when the cohort was divided into male and female subsets after p-value correction (Figure S6D). Geometric mean scores for androgenic steroids and significant SPEAR analytes (which contained several top metabolites from the androgenic steroids pathway) were significantly associated with LC only in males (Figure S6D). The androgenic steroid scores were higher in males than females irrespective of LC status (Figure S6D), corresponding to the higher overall recovery factor score in males (Figure S6B). Notably, the combined score of the top 73 unique features from Hallmark Heme Metabolism, androgenic steroids, and significant SPEAR analytes remained significantly associated with LC in both sexes (Figure S6D).

### Vaccination is not associated with altered recovery factor scores

Our cohort was enrolled prior to the national rollout of SARS-CoV-2 vaccines for the general population. During the longitudinal post-hospitalization follow-up period, as vaccines became broadly available, close to 75% of the participants in the IMPACC Convalescent cohort received a SARS-CoV-2 vaccination (Figure S7A-B). To assess the potential influence of the vaccine response on the immune profiling data and thus the recovery factor, we compared recovery factor scores per visit for events occurring before and after the first vaccination dose, as well as events occurring within a three-week period after any vaccination dose, when vaccine responses have been shown to impact immune profiles^72,73^. No significant difference was found in recovery factor scores across these comparisons, indicating a negligible effect of vaccination on immune profiles related to LC in our patient cohort (Figure S7 C-D).

### Recovery factor scores during the acute disease phase associate with LC status during convalescence

We next investigated whether the immune elements identified in the recovery factor were detectable during the acute infection phase, prior to development of LC. We computed recovery factor scores using immune profiling data from all participants in the Convalescent cohort during their acute phase visits (Visits 1 to 6, spanning hospital admission through 26-35 days post-admission). Remarkably, we found that recovery factor scores were significantly higher in MIN versus LC participants as early as hospital admission (Visit 1) and consistently during the acute period (Figure 5A; Figure S8A). Recovery factor scores were also significantly higher in MIN versus COG groups and MIN versus PHY groups in the acute phase when assessed across the 28-day time course (Figure 5B; Figure S8B). Geometric means of heme metabolism and androgenic steroid pathway analytes from the recovery factor, as well as the 26 significant SPEAR recovery factor analytes were also significantly associated with LC status during the acute phase. The combined geometric mean score of analytes from these three feature groups in acute phase data associated most significantly with MIN versus LC status (Figure 5C), as it did previously in the convalescent phase (Figure 3C).

**Figure 5.**
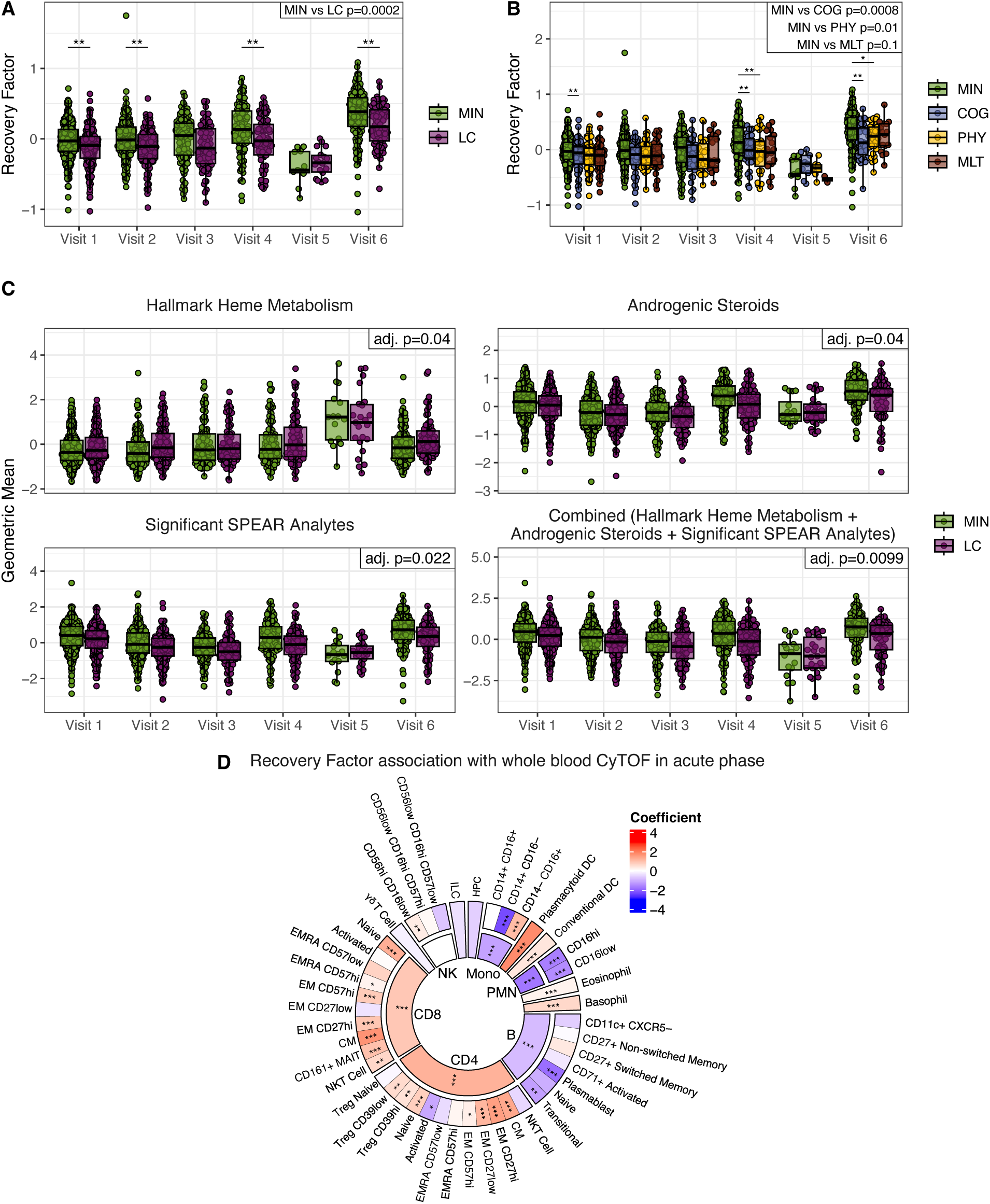
Recovery factor scores in acute phase data associate with eventual LC status. **(A)** Recovery factor scores during the acute disease phase for participants in the LC and MIN groups within 72h of hospital admission (Visit 1) and at day 4 (Visit 2), day 7 (Visit 3), day 14 (Visit 4), day 21 (Visit 5), and day 28 (Visit 6) after admission. **(B)** Recovery factor scores during the acute disease phase for participants in individual PRO clusters. **(C)** Geometric mean of analytes in enriched gene and metabolic sets and/or significant SPEAR analytes during the acute phase. No individual per-visit comparisons were significant after p-value correction. P-values in top-right box in A-C show the significance of the recovery factor score or geometric mean signature association with MIN vs LC or pairwise PRO cluster combinations. Bars above the boxplots show the pairwise significance across groups in a per-visit comparison (** p<0.01, *p<0.05). **(D)** Recovery factor scores association with whole blood CyTOF immune cell populations during the acute phase (* adj. p-value < 0.05, ** adj. p-value < 0.01, *** adj. p-value < 0.001). See also Figure S8.

We further assessed whether altered circulating immune cell composition in the acute phase of disease could contribute to acute-phase recovery factor scores. Association testing of recovery factor scores with whole blood CyTOF measurements during the acute disease phase showed that CD4+ and CD8+ T cells, conventional and plasmacytoid dendritic cells, eosinophils, basophils, and CD56hi CD16low natural killer (NK) cells were significantly positively associated with the recovery factor scores. Within the CD4+ T cell compartment, naïve, central memory (TCM), and effector memory (TEM) subsets, as well as non-naive regulatory T cells (Treg) were significantly associated with recovery factor scores, while activated CD4+ T cells were inversely correlated. Within the CD8+ T cell compartment, naïve, TCM, and TEM subsets were positively associated with recovery factor scores, as were NKT cells and CD161+ MAIT cells. In contrast, monocytes, neutrophils, B cells, and plasmablasts in the acute phase were negatively associated with recovery factor scores (Figure 5D). These findings are consistent with a previous study that found higher plasmablast counts and lower total CD4+ T, total CD8+ T, CD4+ TEM, CD8 TEM, Treg, NK, and dendritic cell counts in immune-cell populations sampled at days 0-14 after infection in COVID-19 patients who experienced persisting symptoms at days 91-180 after infection^29^. The similarities across both studies are indicative of an acute blood immune cell type signature of LC that is robust to variance in patient cohorts and LC definition.

In summary, our findings indicate that the major biologic signatures of the recovery factor that stratify LC from recovered participants in the convalescent phase – elevated heme metabolism gene signatures, reduced androgenic steroids, increased circulating inflammatory mediators, and increased monocytes and neutrophils – are evident early in the acute phase of disease.

### Acute-phase recovery factor scores distinguish acute disease severities and predict LC risk irrespective of acute severity

We investigated the full IMPACC study cohort (n = 1,148 participants with at least one sample measurement for the omics modalities included in our model) to assess whether recovery factor scores determined from acute phase data would associate with patient severity trajectory group assignments. For this analysis we included participants who did not survive beyond 28 days post-hospital admission and participants without biospecimens and/or surveys during the convalescent phase. Recovery factor scores were significantly associated longitudinally with acute disease trajectory groups and were highest in participants with milder disease courses (TG1-TG3) and lowest in participants with the most severe acute disease trajectories (TG4 and TG5) (Figure 6A). Acute-phase recovery factor scores increased over time for participants in all trajectory groups except TG5, the most severe group in which participants died by day 28 after hospital admission (Figure 6A). To assess whether the association between acute recovery factor scores and convalescent LC status was simply due to acute recovery factor scores being an indicator of acute disease severity, we repeated the association test including trajectory group as a covariate at each visit (Figure 6B) and longitudinally (Figure S8C). LC status was still significantly associated with acute recovery factor scores even after taking trajectory group into account. These findings suggest that recovery factor scores in the acute phase contain valuable information for predicting convalescent LC status beyond its correlation with acute disease severity.

**Figure 6.**
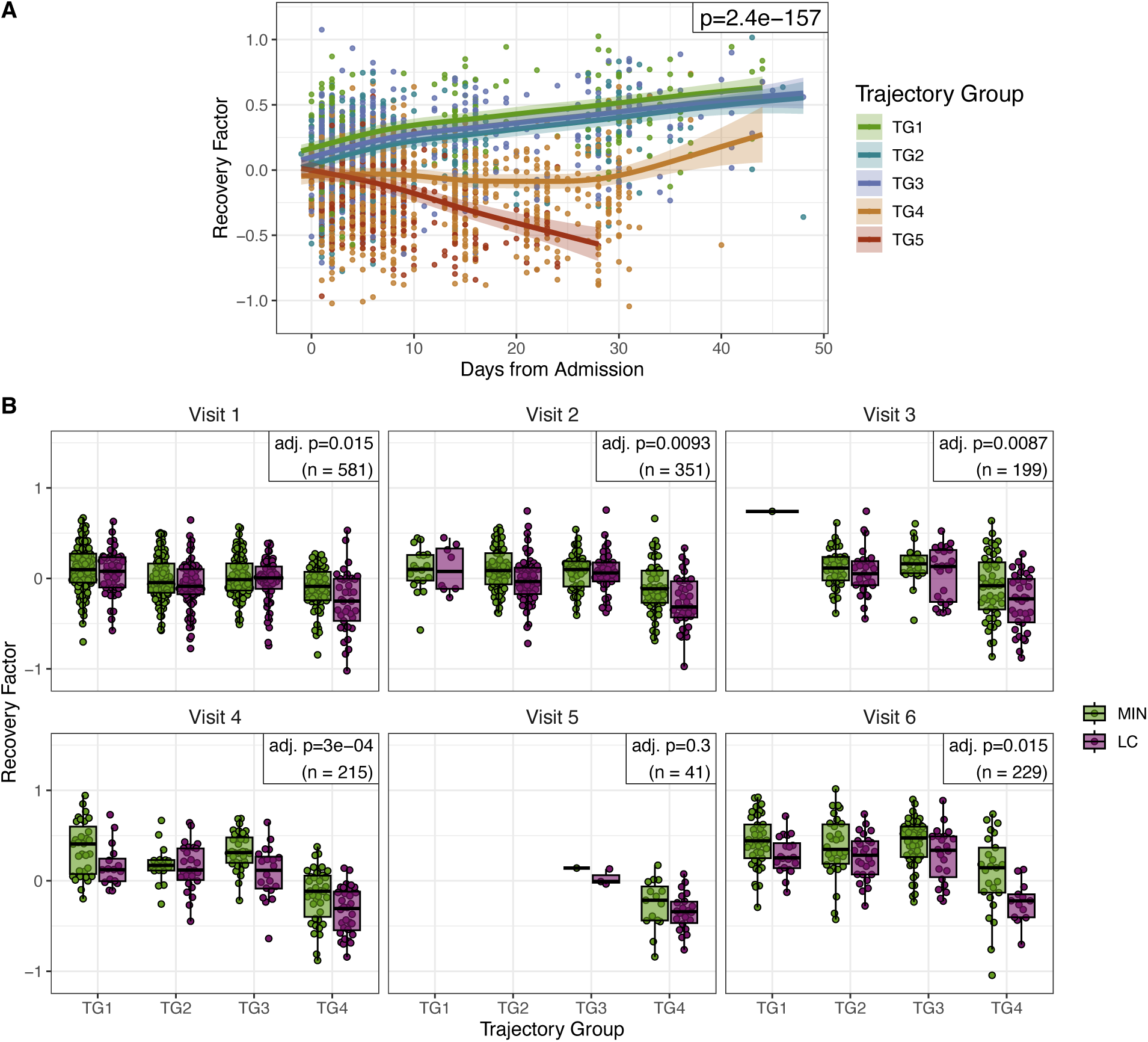
Recovery factor scores associate with acute disease phase trajectory groups, but identify LC irrespective of acute severity. **(A)** Longitudinal analysis of acute recovery factor scores for the full IMPACC cohort stratified by trajectory group (N=1,148 participants). P-value shows the significance of the trajectory group term in a longitudinal model correcting for age and sex as fixed effects and enrollment site and participant ID as random effects. **(B)** Recovery factor scores in the acute phase by convalescent MIN/LC label, stratified by acute trajectory group and visit number. P-values show significance in distinguishing MIN vs. LC labels in linear mixed models with sex, discretized admit age, and trajectory group as fixed effects and enrollment site and participant ID as random effects, performed separately for each acute visit and corrected across all visits.

### Machine learning models based on the recovery factor scores at the acute or the convalescent phase together with clinical features predict LC status during the convalescent phase

Building on the strength of the recovery factor to predict LC even from the acute phase of disease, we assessed whether a combination of the recovery factor scores and clinical characteristics could improve the predictive performance. We selected clinical features including the routinely recorded elements of age at enrollment, sex, body mass index (BMI), length of hospital stay, Sequential Organ Failure Assessment (SOFA) score, along with Spike IgG antibody titers and viral load (SARS-CoV-2 N1 PCR) during the acute phase of disease, which were previously associated with LC status in the IMPACC cohort^18^ (Figure S9), as well as the presence of comorbidities including hypertension, diabetes, chronic cardiac disease, chronic kidney disease, malignant neoplasms, chronic neurological disorders, liver disfunction/failure, history of transplants, smoking/vaping, asthma, respiratory diseases other than asthma, substance use, HIV infection, and the total number of comorbidities (Figure S10A). Lasso models trained exclusively on the mean recovery factor scores during the acute phase (SPEAR acute, mean cross-validation AUROC 0.64) outperformed models trained exclusively on clinical features (Clinical, mean cross-validation AUROC 0.63) (Figure S10B). The performance in predicting LC was further improved when training models on the combination of baseline and acute-phase clinical features and recovery factor scores (SPEAR acute + clinical, mean cross-validation AUROC 0.66) (Figure S10B). The lasso model trained on the recovery factor scores during the convalescent phase (mean cross-validation AUROC 0.74) also outperformed the clinical-only model and was slightly improved when combined with clinical features (mean cross-validation AUROC 0.75) (Figure S10C). In addition, we assessed whether a reduced set of analytes based on the 26 SPEAR significant analytes could achieve similar predictive performance to the models trained on the full recovery factor (Figure S10C). Such streamlined models would be advantageous in a clinical setting or to develop clinical diagnostics tests to identify individuals with LC more easily. Even though the best performing reduced analyte set models (SPEAR significant analytes) had a lower predictive power than the models trained on the full convalescent recovery factor scores, this sparse model achieved a mean cross-validation AUROC of 0.69, suggesting its utility.

We further evaluated whether single time point measurements of the recovery factor scores could achieve similar performance as the models with averaged scores over the acute or convalescent phases. Interestingly, we observed a slowly increasing trend in predictive performance for each individual time point with increased visit numbers during earlier visits, which peaked at Visit 8 (6 months after hospital discharge) with a mean cross-validation AUROC of 0.74 (Figure S10D). The 6-month time point was also the strongest individual time point predictor when the model included the clinical features (mean cross-validation AUROC of 0.76) (Figure S10D). Notably, recovery factor scores as early as Visit 1 provided predictive performance (mean cross-validation AUROC 0.63), indicating that the recovery factor captures early predictive features of LC during the acute disease phase, albeit the signal at this early time point is not as strongly predictive as later in the convalescent phase.

## DISCUSSION

In this study, we applied supervised multi-omics integration methods to identify biologic features associated with LC in 513 participants from the IMPACC cohort. We took advantage of data availability from this cohort that was followed longitudinally after hospitalization with COVID-19 through the first 28 days of the acute phase of disease and subsequently for up to one year after hospital discharge^18,70^. The IMPACC cohort is unique in its comprehensive inclusion of clinical data, biospecimens, and quarterly patient-reported outcome surveys, combined with multi-omics immunophenotyping at multiple time points throughout the acute and convalescent phases of the disease. A previous study of this cohort identified demographic and clinical risk factors associated with the development of LC, which included female sex, comorbidities such as chronic heart, lung, and neurologic diseases, and a longer length of hospital stay^18^. Here, we report the first study of the IMPACC cohort to analyze biologic data collected during the convalescent phase of disease, allowing us to identify a multi-omics “recovery factor” capable of discriminating participants who recover with minimal deficits from those who experience a variety of clinical LC symptoms. Notably, we find that as early as 72h after hospital admission for COVID-19, recovery factor scores predict which patients, irrespective of their acute disease severity, will go on to experience LC. Biologic features associated with the recovery factor score indicate that reduced androgenic steroid levels, increased heme metabolism signatures, and persistent elevation of inflammation-associated serum proteins are hallmarks of LC that both identify individuals with LC during convalescence and predict which acute COVID-19 patients will experience LC.

Increased levels of androgenic steroids in serum were positively associated with recovery factor scores and with participants who recovered from COVID-19 with minimal deficits. Seven of the twelve leading-edge androgenic steroid metabolites, as well as Oxysterol Binding Protein 2 (OSBP2), a leading-edge gene in the Hallmark Heme Metabolism pathway, were also included within the list of 26 analytes that were statistically significant within the recovery factor. Limited studies have elucidated the role of reduced androgenic steroids in LC, but in agreement with our findings, lower testosterone levels have been associated with increased LC symptomatology in both males and females^31^. We observed that several intermediate metabolites in the canonical steroid hormone biosynthesis pathway were associated with the recovery factor and were decreased in LC participants, including sulfated forms of testosterone precursors (pregnenolone and DHEA) and downstream metabolites (androsterone, epiandrosterone, and 5alpha-androstan-3beta,17beta-diol). Notably, our list of top androgenic steroid metabolites has strong overlap with the androgenic steroid signature from an independent all-female cohort of healthy controls compared to myalgic encephalomyelitis / chronic fatigue syndrome (ME/CFS) patients^74^. All six androgenic steroid metabolites that were significantly elevated in healthy controls in this study (DHEA-S, androstenediol (3alpha, 17alpha) monosulfate (2), androstenediol (3beta,17beta) disulfate (2), 5alpha-androstan-3beta,17alpha-diol disulfate, androsterone sulfate, epiandrosterone sulfate)^74^ were also statistically significant in our recovery factor, strongly implicating this signature in the shared symptomology, such as fatigue, post exertional malaise, and sleep disturbances, between ME/CFS and LC^75^. In our study, a geometric mean score consisting of leading-edge metabolites from the androgenic steroids pathway significantly differentiated MIN vs. LC in the entire cohort. Nonetheless, we note that when the cohort was subset into males and females, the geometric mean scores for the androgenic steroid pathway only significantly stratified LC from recovered male participants, although levels of androgenic steroid scores also trended lower in females with LC.

The fact that pregnenolone (sulfate) and many of its downstream metabolites were all significantly elevated in the recovery factor, despite being generated by distinct enzymes, suggests that a rate-limiting enzymatic step upstream of pregnenolone may explain the observed signature. We observed that cholesterol, which is cleaved by the mitochondrial enzyme CYP11A1 to produce pregnenolone in the first step of the steroid hormone biosynthesis process, was detected in the metabolomics assay but was not identified as a significant metabolite in the recovery factor. CYP11A1 was not detected in our proteomics or transcriptomic assays, likely due to localized activity in the adrenal gland. Further experimentation is needed to test if impaired generation of pregnenolone from cholesterol in LC is responsible for decreased levels of intermediate androgenic steroid metabolites. Alternatively, the observed decrease of sulfated androgenic steroid levels in LC may reflect altered activity of sulfatases or sulfotransferase enzymes. Notably, some sulfonated steroids, such as DHEAS, a leading-edge gene in the androgenic steroid pathway and a top significant analyte in the SPEAR recovery factor, have been shown to exhibit immunosuppressive and anti-inflammatory effects, particularly in neuroinflammation^76^. In addition, testosterone has shown to play an immunomodulatory role and is often reduced in patients with other critical illnesses^77^. Thus, elevated sulfated steroids in the recovery factor may contribute to the reduction in inflammation associated with recovery from COVID-19 with minimal deficits^76^.

In addition to altered androgenic steroids, the transcriptional signature of heme metabolism in PBMCs was inversely associated with the recovery factor, such that LC participants had elevated expression of genes associated with the heme metabolism pathway during the convalescent period. As with androgenic steroids, we found the leading-edge genes of the heme metabolism pathway were also higher during the acute phase of disease in participants who would later experience LC. Notably, overexpression of a heme metabolism signature in blood was recently reported for a separate cohort of 102 participants, including non-hospitalized and hospitalized COVID-19 patients evaluated 1-3 months after infection, which found this pathway to be enriched in participants experiencing persisting symptoms^29^. The authors related the elevated heme metabolism signature to stress erythropoiesis induced in response to inflammation-associated anemia driven by IL-6-mediated hepcidin upregulation^29,78^. In that study, participants who experienced LC had reduced iron and HGB 2 weeks to 1 month after COVID-19 infection. This iron restriction was proposed not only to induce anemia, but also to impair lymphocyte function. The subsequent delayed resolution of acute infection might result in sustained inflammation and persistent anemia up to at least 6 months after infection, which may partially explain the systemic symptomatology of acute COVID-19 and LC. Anemia of inflammation, also known as anemia of chronic disease, is a common complication associated with chronic inflammatory illnesses such as chronic kidney disease, congestive heart failure, as well as ICU admission^79^. Anemia as a complication was also significantly negatively associated with the recovery factor and was shown to be associated with PRO clusters in our previous study^18^, whereas hemoglobin and hematocrit baseline measurements showed a significant positive association with the recovery factor. Our findings are largely consistent with this study, as we also find evidence of persistent inflammation in LC participants: inflammation-associated serum factors, such as CXCL9, CSF1, and FGF21, were identified by SPEAR as significant analytes in the recovery factor, albeit they did not reach significance when associated individually with LC status. Of note, FGF21 measured in the acute phase was previously associated with cognitive and multidomain deficit PRO clusters relative to MIN in this cohort^18^, and has been proposed as a biomarker for chronic inflammation in myalgic encephalomyelitis/chronic fatigue syndrome (ME/CFS)^80^, a complex chronic disease that overlaps clinically with LC. Furthermore, LRG1, which is activated by the inflammatory IL-6/STAT3 pathway is significantly elevated in LC participants, perhaps contributing to vascular pathology in LC^60–62^. Although IL-6 (SPEAR Bayesian posterior selection probability = 0.86) did not reach significance in the SPEAR factor, it was within the top 50 analytes of the factor. Along with these persistent signatures of inflammation, we also identified elevated heme metabolism gene expression signatures as a key feature of LC. In fact, expression levels of the leading-edge heme metabolism genes we identified in our study could also predict LC participants in another cohort^29^, demonstrating that the same heme metabolism signature during the acute phase of COVID-19 can predict which patients will experience LC when tested in participants from distinct studies. Together, these findings point to persistent inflammation, driving anemia and stress erythropoiesis as key drivers of LC.

Across all time points, consistent immune cell subsets were associated with the recovery factor. In both the acute and convalescent disease phases, CD161+ MAIT cell frequencies were positively associated with the recovery factor, while monocyte, neutrophil, and CD14+ CD16-classical monocyte cell frequencies were negatively associated with the recovery factor. However, the B cell population is negatively associated with the recovery factor during the acute disease phase, but becomes positively associated during convalescence. We also found several points of agreement for immune cell frequencies in the recovery factor with previous reports of LC subjects. While comparing immune composition across cohorts can be challenging due to differences in cohort construction and cell type annotation, we found that CD4+ and CD8+ T cells, plasmacytoid DCs, and conventional DCs are positively associated with the recovery factor during the acute phase of disease, while monocytes, neutrophils, and B cells are negatively associated, suggesting reduced T cell immunity relative to inflammatory innate immunity in the acute phase of disease in individuals most susceptible to LC. This finding is in keeping with cellular trends observed in Hanson et al.^29^ in the acute phase of disease and in males during convalescence by Silva and colleagues^31^. Consistent with Klein et al.^26^, we do not find significant associations of the recovery score with naive CD4+ or naive CD8+ T cell subsets in the convalescent phase of COVID-19.

A previous study from IMPACC identified a ‘severity factor’ that significantly associated with clinical outcomes during the acute phase of disease^39^. Given that both the severity factor and the recovery factor are associated with inflammatory signatures during acute COVID-19, we compared immune cell types associated with the recovery factor to those associated with the severity factor at this stage of disease. There were notable similarities: for example, the combined monocyte, B cell, and neutrophil populations are negatively associated with the recovery factor and positively associated with the severity factor. Likewise, CD4+ T cell and CD8+ T cell populations are positively associated with the recovery factor and negatively associated with the severity factor. These cellular associations suggest that inefficient adaptive immunity during the acute phase of disease, with elevated frequencies of inflammatory innate cells contribute to LC susceptibility. This model is consistent with our past^18^ and current findings in the IMPACC cohort that reduced levels of anti-SARS-CoV-2 antibodies and increased viral titers in patients within the first 72 hours of hospital admission are associated with participants who will experience LC (Figure S9). These findings are also consistent with other reports that hospitalized individuals are more susceptible to LC than non-hospitalized individuals^14,81^. Nonetheless, our recovery factor predicts which patients will experience LC, irrespective of acute disease severity, indicating that the model has learned features of COVID-19 beyond inflammation that are associated with COVID-19 severity.

Although symptom groups, such as respiratory symptoms, were found to be significantly associated with LC in this cohort^18^, the multi-omics recovery factor does not associate with a particular clinical symptom group. Instead, it captures biomarkers predictive of global physical deficits, as reported by patients after acute COVID-19 disease. While assessing the entire multi-omics SPEAR factor in convalescent patients is impractical, our findings indicate that assessing the 26 significant SPEAR analytes would aid in LC diagnoses (Fig S10c).

This study has several limitations. The reliance on self-reported survey data to identify symptoms and classify participants into MIN/LC and individual PRO clusters may introduce potential biases. To address this, population-normalized PRO measure scores were utilized, and comparisons were made to pre-illness health status when possible. Additionally, the surveys were developed early in the pandemic (March 2020), before the full spectrum of LC symptoms was characterized, and thus did not capture current commonly-recognized manifestations such as brain fog, fatigue, sleep disturbances, neuropathy, and dysautonomia. Self-selection bias may also be present, as patients with severe LC symptoms might have been less likely to respond to the surveys. As the study cohort was recruited during early phases of the pandemic (May 2020 through March 2021), it consists of individuals infected with the original SARS-CoV-2 strain and does not include data on subsequent variants of concern, limiting the generalization of the findings to other later dominant strains. Vaccination data were self-reported and limited to the post-acute phase since enrollment was largely completed prior to vaccine rollout, and exact vaccination dates were unavailable for some participants. Latent virus reactivation was not considered in this model. Furthermore, as part of the study design, all participants in the IMPACC cohort were hospitalized for COVID-19. Consequently, the multi-omics factors were constructed without incorporating profiles from individuals with COVID-19 who did not require hospitalization or healthy controls, potentially introducing a bias toward those with more severe disease. However, we were able to discover in our hospitalized cohort molecular signatures that were previously observed in cohorts that included non-hospitalized and asymptomatic SARS-CoV-2 infected participants^29^.

Despite these limitations, the study possesses significant strengths. In addition to the prospective design, with acute and convalescent longitudinal multi-omics profiling, enrollment through multiple sites across the United States enhances the diversity and broad representation of the cohort and mitigates potential participant recruitment biases, contributing to the robustness of the findings.

In conclusion, supervised multi-omics factor construction of immune profiling data from SARS-CoV-2 infected participants who recovered with minimal deficits or experienced LC indicates that immune cell types and serum factors associated with inflammation, reduced androgenic steroids, and an elevated heme metabolism signature predict which participants will experience LC, irrespective of acute disease severity. Moreover, these signatures are maintained into convalescence, indicating that persistent inflammation is likely a key driver of LC. Further studies will be needed to determine why inflammation persists in some COVID-19 patients. We did not assess persistent SARS-CoV-2 viral loads or viral reactivation in this study; however, a recent study of IMPACC data^82^ suggests reactivation of latent viruses could contribute, consistent with other studies^26,30,31^. Altogether, our data, paired with prior congruent reports, suggest that impaired lymphocyte function early in COVID-19 reduces cellular and humoral adaptive immunity and contributes to high SARS-CoV-2 viral loads. Elevated viral loads can trigger innate immune cell responses that increase inflammatory cytokines, driving inflammation-associated anemia that further reduces lymphocyte function, which could enable reactivation of latent viruses. Such persistent inflammation that is not successfully resolved likely leads to LC pathology. Strategies to break the cycle of inflammation and correct the inflammation-associated anemia may promote recovery from LC, and merit further investigation.

## Supporting information

Gabernet Supplemental Figures

Gabernet Supplemental Methods

## RESOURCE AVAILABILTY

### Lead Contact

Requests for further information should be directed to Lauren Ehrlich

### Materials availability

This study did not generate new, unique reagents.

### Data and Code availability

Data used in this study is available on the ImmPort repository under accession number SDY1760 and in the Database of Genotypes and Phenotypes (dbGaP) under the accession number phs002686.v2.p2.

All code used in this study is deposited on Bitbucket (https://bitbucket.org/kleinstein/impacc-public-code/src/master/multiomics-longcovid/).

## CONSORTIA

The members of the IMPACC Network are:

**Clinical and Data Coordinating Center (CDCC), Precision Vaccines Program, Boston Children’s Hospital, Harvard Medical School, Boston, MA 02115, USA:**

Al Ozonoff, Joann Diray-Arce, Jing Chen, Alvin T. Kho, Carly E. Milliren, Annmarie Hoch, Ana C. Chang, Kerry McEnaney, Caitlin Syphurs, Brenda Barton, Claudia Lentucci, Maimouna D. Murphy, Mehmet Saluvan, Tanzia Shaheen, Shanshan Liu, Marisa Albert, Arash Nemati Hayati, Robert Bryant, James Abraham, Mitchell Cooney, Meagan Karoly

**Benaroya Research Institute, University of Washington, Seattle, WA 98101, USA:**

Matthew C. Altman, Naresh Doni Jayavelu, Scott Presnell, Bernard Kohr, Tomasz Jancsyk, Azlann Arnett

**La Jolla Institute for Immunology, La Jolla, CA 92037, USA:**

Bjoern Peters, James A. Overton, Randi Vita, Kerstin Westendorf

**Knocean Inc. Toronto, ON M6P 2T3, Canada:**

James A. Overton

**Precision Vaccines Program, Boston Children’s Hospital, Harvard Medical School, Boston, MA 02115, USA:**

Ofer Levy, Hanno Steen, Patrick van Zalm, Benoit Fatou, Kinga K. Smolen, Arthur Viode, Simon van Haren, Meenakshi Jha, David Stevenson, Sanya Thomas, Boryana Petrova, Naama Kanarek

**Brigham and Women’s Hospital, Harvard Medical School, Boston, MA 02115, USA:**

Lindsey R. Baden, Kevin Mendez, Jessica Lasky-Su, Alexandra Tong, Rebecca Rooks, Michael Desjardins, Amy C. Sherman, Stephen R. Walsh, Xhoi Mitre, Jessica Cauley, Xiaofang Li, Bethany Evans, Christina Montesano, Jose Humberto Licona, Jonathan Krauss, Nicholas C. Issa, Jun Bai Park Chang, Natalie Izaguirre

**Metabolon Inc, Morrisville, NC 27560, USA:**

Scott R. Hutton, Greg Michelotti, Kari Wong

**Prevention of Organ Failure (PROOF) Centre of Excellence, University of British Columbia, Vancouver, BC V6T 1Z3, Canada:**

Scott J. Tebbutt, Casey P. Shannon

**Case Western Reserve University and University Hospitals of Cleveland, Cleveland, OH 44106, USA:**

Rafick-Pierre Sekaly, Slim Fourati, Grace A. McComsey, Paul Harris, Scott Sieg, George Yendewa, Mary Consolo, Heather Tribout, Susan Pereira Ribeiro

**Drexel University, Tower Health Hospital, Philadelphia, PA 19104, USA:**

Charles B. Cairns, Elias K. Haddad, Michele A. Kutzler, Mariana Bernui, Gina Cusimano, Jennifer Connors, Kyra Woloszczuk, David Joyner, Carolyn Edwards, Edward Lee, Edward Lin, Nataliya Melnyk, Debra L. Powell, James N. Kim, I. Michael Goonewardene, Brent Simmons, Cecilia M. Smith, Mark Martens, Brett Croen, Nicholas C. Semenza, Mathew R. Bell, Sara Furukawa, Renee McLin, George P. Tegos, Brandon Rogowski, Nathan Mege, Kristen Ulring, Pam Schearer, Judie Sheidy, Crystal Nagle

**MyOwnMed Inc., Bethesda, MD 20817, USA:**

Vicki Seyfert-Margolis

**Emory School of Medicine, Atlanta, GA 30322, USA:**

Nadine Rouphael, Steven E. Bosinger, Arun K. Boddapati, Greg K. Tharp, Kathryn L. Pellegrini, Brandi Johnson, Bernadine Panganiban, Christopher Huerta, Evan J. Anderson, Hady Samaha, Jonathan E. Sevransky, Laurel Bristow, Elizabeth Beagle, David Cowan, Sydney Hamilton, Thomas Hodder, Amer Bechnak, Andrew Cheng, Aneesh Mehta, Caroline R. Ciric, Christine Spainhour, Erin Carter, Erin M. Scherer, Jacob Usher, Kieffer Hellmeister, Laila Hussaini, Lauren Hewitt, Nina Mcnair, Susan Pereira Ribeiro, Sonia Wimalasena

**Icahn School of Medicine at Mount Sinai, New York, NY 10029, USA:**

Ana Fernandez-Sesma, Viviana Simon, Florian Krammer, Harm Van Bakel, Seunghee Kim-Schulze, Ana Silvia Gonzalez-Reiche, Jingjing Qi, Brian Lee, Juan Manuel Carreño, Gagandeep Singh, Ariel Raskin, Johnstone Tcheou, Zain Khalil, Adriana van de Guchte, Keith Farrugia, Zenab Khan, Geoffrey Kelly, Komal Srivastava, Lily Q. Eaker, Maria C. Bermúdez-González, Lubbertus C.F. Mulder, Katherine F. Beach, Miti Saksena, Deena Altman, Erna Kojic, Levy A. Sominsky, Arman Azad, Dominika Bielak, Hisaaki Kawabata, Temima Yellin, Miriam Fried, Leeba Sullivan, Sara Morris, Giulio Kleiner, Daniel Stadlbauer, Jayeeta Dutta, Hui Xie, Manishkumar Patel, Kai Nie, Brian Monahan

**Immunai Inc., New York, NY 10016, USA:**

Adeeb Rahman

**Oregon Health & Science University, Portland, OR 97239, USA:**

William B. Messer, Catherine L. Hough, Sarah A.R. Siegel, Peter E. Sullivan, Zhengchun Lu, Amanda E. Brunton, Matthew Strand, Zoe L. Lyski, Felicity J. Coulter, Courtney Micheletti

**Stanford University School of Medicine, Palo Alto, CA 94305, USA:**

Holden Maecker, Bali Pulendran, Kari C. Nadeau, Yael Rosenberg-Hasson, Michael Leipold, Natalia Sigal, Angela Rogers, Andrea Fernandes, Monali Manohar, Evan Do, Iris Chang, Alexandra S. Lee, Catherine Blish, Henna Naz Din, Jonasel Roque, Linda N. Geng, Maja Artandi, Mark M. Davis, Neera Ahuja, Samuel S. Yang, Sharon Chinthrajah, Thomas Hagan, Tyson H. Holmes, Koji Abe

**David Geffen School of Medicine at the University of California Los Angeles, Los Angeles CA 90095, USA:**

Elaine F. Reed, Joanna Schaenman, Ramin Salehi-Rad, Adreanne M. Rivera, Harry C. Pickering, Subha Sen, David Elashoff, Dawn C. Ward, Jenny Brook, Estefania Ramires-Sanchez, Megan Llamas, Claudia Perdomo, Clara E. Magyar, Jennifer Fulcher

**University of California San Francisco, San Francisco, CA 94115, USA:**

David J. Erle, Carolyn S. Calfee, Carolyn M. Hendrickson, Kirsten N. Kangelaris, Viet Nguyen, Deanna Lee, Suzanna Chak, Rajani Ghale, Ana Gonzalez, Alejandra Jauregui, Carolyn Leroux, Luz Torres Altamirano, Ahmad Sadeed Rashid, Andrew Willmore, Prescott G. Woodruff, Matthew F. Krummel, Sidney Carrillo, Alyssa Ward, Charles R. Langelier, Ravi Patel, Michael Wilson, Ravi Dandekar, Bonny Alvarenga, Jayant Rajan, Walter Eckalbar, Andrew W. Schroeder, Gabriela K. Fragiadakis, Alexandra Tsitsiklis, Eran Mick, Yanedth Sanchez Guerrero, Christina Love, Lenka Maliskova, Michael Adkisson, Aleksandra Leligdowicz, Alexander Beagle, Arjun Rao, Austin Sigman, Bushra Samad, Cindy Curiel, Cole Shaw, Gayelan Tietje-Ulrich, Jeff Milush, Jonathan Singer, Joshua J. Vasquez, Kevin Tang, Legna Betancourt, Lekshmi Santhosh, Logan Pierce, Maria Tecero Paz, Michael Matthay, Neeta Thakur, Nicklaus Rodriguez, Nicole Sutter, Norman Jones, Pratik Sinha, Priya Prasad, Raphael Lota, Saurabh Asthana, Sharvari Bhide, Tasha Lea, Yumiko Abe-Jones

**Yale School of Medicine, New Haven, CT 06510, USA:**

David A. Hafler, Ruth R. Montgomery, Albert C. Shaw, Steven H. Kleinstein, Jeremy P. Gygi, Dylan Duchen, Shrikant Pawar, Anna Konstorum, Ernie Chen, Chris Cotsapas, Xiaomei Wang, Charles Dela Cruz, Akiko Iwasaki, Subhasis Mohanty, Allison Nelson, Yujiao Zhao, Shelli Farhadian, Hiromitsu Asashima, Omkar Chaudhary, Andreas Coppi, John Fournier, M. Catherine Muenker, Khadir Raddassi, Michael Rainone, William Ruff, Syim Salahuddin, Wade L. Shulz, Pavithra Vijayakumar, Haowei Wang, Esio Wunder Jr., H. Patrick Young, Albert I. Ko, Gisela Gabernet

**Yale School of Public Health, New Haven, CT 06510, USA:**

Denise Esserman, Leying Guan, Anderson Brito, Jessica Rothman, Nathan D. Grubaugh, Kexin Wang, Leqi Xu

**Baylor College of Medicine and the Center for Translational Research on Inflammatory Diseases, Houston, TX 77030, USA:**

David B. Corry, Farrah Kheradmand, Li-Zhen Song, Ebony Nelson

**Oklahoma University Health Sciences Center, Oklahoma City, OK 73104, USA:**

Jordan P. Metcalf, Nelson I. Agudelo Higuita, Lauren A. Sinko, J. Leland Booth, Douglas A. Drevets, Brent R. Brown

**University of Arizona, Tucson AZ 85721, USA:**

Monica Kraft, Chris Bime, Jarrod Mosier, Heidi Erickson, Ron Schunk, Hiroki Kimura, Michelle Conway, Dave Francisco, Allyson Molzahn, Connie Cathleen Wilson, Ron Schunk, Trina Hughes, Bianca Sierra

**University of Florida, Gainesville, FL 32611, USA:**

Mark A. Atkinson, Scott C. Brakenridge, Ricardo F. Ungaro, Brittany Roth Manning, Lyle Moldawer

**University of Florida, Jacksonville, FL 32218, USA:**

Jordan Oberhaus, Faheem W. Guirgis

**University of South Florida, Tampa FL 33620, USA:**

Brittney Borresen, Matthew L. Anderson

**The University of Texas at Austin, Austin, TX 78712, USA:**

Lauren I. R. Ehrlich, Esther Melamed, Cole Maguire, Dennis Wylie, Justin F. Rousseau, Kerin C. Hurley, Janelle N. Geltman, Nadia Siles, Jacob E. Rogers, Pablo Guaman Tipan

## ACKNOWLEDGMENTS

This research was supported by the following grants: NIH (3U01AI167892-03S2, 3U01AI167892-01S2, 5R01AI135803-03, 5U19AI118608-04, 5U19AI128910-04, 4U19AI090023-11, 4U19AI118610-06, R01AI145835-01A1S1, 5U19AI062629-17, 5U19AI057229-17, 5U19AI057229-18, 5U19AI125357-05, 5U19AI128913-03, 3U19AI077439-13, 5U54AI142766-03, 5R01AI104870-07, 3U19AI089992-09, 3U19AI128913-03, and 5T32DA018926-18); NIAID, NIH (3U19AI1289130, U19AI128913-04S1, and R01AI122220); NCATS, NIH UM1TR004528 and National Science Foundation (DMS2310836).

We thank the participants of the study for their voluntary enrollment and contribution of samples for this work. See supplement information for details on the IMPACC Network. We acknowledge the assistance of the following individuals: Tyson H. Holmes (Human Immune Monitoring Center, Stanford University), Sanya Thomas, Mitchell Cooney, Shun Rao, Sofia Vignolo, and Elena Morrocchi (all from the CDCC); Arash Naeim, Marianne Bernardo, Sarahmay Sanchez, Shannon Intluxay, Clara Magyar, Jenny Brook, Estefania Ramires-Sanchez, Megan Llamas, Claudia Perdomo, Clara E. Magyar, and Jennifer A. Fulcher (all from the David Geffen School of Medicine at UCLA); members of the UCLA Center for Pathology Research Services and the Pathology Research Portal; M. Catherine Muenker, Dimitri Duvilaire, Maxine Kuang, William Ruff, Khadir Raddassi, Denise Shepherd, Haowei Wang, Omkar Chaudhary, Syim Salahuddin, John Fournier, Michael Rainone, and Maxine Kuang (all from the Yale School of Medicine). We thank the leadership of Boston Children’s Hospital including Drs. Wendy Chung, Gary Fleisher and Kevin Churchwell for their support for the Precision Vaccines Program.

## DECLARATION OF INTERESTS

The Icahn School of Medicine at Mount Sinai has filed patent applications relating to SARS-CoV-2 serological assays, NDV-based SARS-CoV-2 vaccines influenza virus vaccines and influenza virus therapeutics which lists Florian Krammer as co-inventor. Mount Sinai has spun out a company, Kantaro, to market serological tests for SARS-CoV-2 and another company, Castlevax, to develop SARS-CoV-2 vaccines. Florian Krammer is co-founder and scientific advisory board member of Castlevax. Florian Krammer has consulted for Merck, Curevac, Seqirus, GSK and Pfizer and is currently consulting for 3rd Rock Ventures, Sanofi, Gritstone and Avimex. The Krammer laboratory is also collaborating with Dynavax on influenza vaccine development and with VIR on influenza virus therapeutics development. Viviana Simon is a co-inventor on a patent filed relating to SARS-CoV-2 serological assays (the “Serology Assays”). Ofer Levy is a named inventor on patents held by Boston Children’s Hospital relating to vaccine adjuvants and human in vitro platforms that model vaccine action. His laboratory has received research support from GlaxoSmithKline (GSK) and Pfizer, and he is a co-founder of and advisor to Ovax, Inc that develops opioid vaccines. Charles Cairns serves as a consultant to bioMerieux and is funded by a grant from Bill & Melinda Gates Foundation. James A Overton is a consultant at Knocean Inc. Jessica Lasky-Su serves as a scientific advisor of Precion Inc. Scott R. Hutton, Greg Michelloti and Kari Wong are employees of Metabolon Inc. Vicki Seyfer-Margolis is a current employee of MyOwnMed. Nadine Rouphael reports grants or contracts with Merck, Sanofi, Pfizer, Vaccine Company and Immorna, and has participated on data safety monitoring boards for Moderna, Sanofi, Seqirus, Pfizer, EMMES, ICON, BARDA, and CyanVan, Imunon Micron. N.R. has also received support for meetings/travel from Sanofi and Moderna and honoraria from Virology Education and Krog consulting. Chris Cotsapas is a current employee of Vesalius Therapeutics. Adeeb Rahman is a current employee of Immunai Inc. Steven Kleinstein is a consultant related to ImmPort data repository for Peraton. Nathan Grabaugh is a consultant for Tempus Labs and the National Basketball Association. Akiko Iwasaki is a consultant for 4BIO, Blue Willow Biologics, Revelar Biotherapeutics, RIGImmune, Xanadu Bio, Paratus Sciences. Monika Kraft receives research funds paid to her institution from NIH, ALA; Sanofi, Astra-Zeneca for work in asthma, serves as a consultant for Astra-Zeneca, Sanofi, Chiesi, GSK for severe asthma; is a co-founder and CMO for RaeSedo, Inc, a company created to develop peptidomimetics for treatment of inflammatory lung disease. Esther Melamed received research funding from Babson Diagnostics and honorarium from Multiple Sclerosis Association of America and has served on the advisory boards of Genentech, Horizon, Teva, and Viela Bio. Carolyn Calfee receives research funding from NIH, FDA, DOD, Roche-Genentech and Quantum Leap Healthcare Collaborative as well as consulting services for Janssen, Vasomune, Gen1e Life Sciences, NGMBio, and Cellenkos. Wade Schulz was an investigator for a research agreement, through Yale University, from the Shenzhen Center for Health Information for work to advance intelligent disease prevention and health promotion; collaborates with the National Center for Cardiovascular Diseases in Beijing; is a technical consultant to Hugo Health, a personal health information platform; cofounder of Refactor Health, an AI-augmented data management platform for health care; and has received grants from Merck and Regeneron Pharmaceutical for research related to COVID-19. Grace A McComsey received research grants from Rehdhill, Cognivue, Pfizer, and Genentech, and served as a research consultant for Gilead, Merck, Viiv/GSK, and Janssen. Linda N. Geng received research funding paid to her institution from Pfizer, Inc. Catherine Hough receives support through her institution from the NIH and CDC. David Hafler has received research funding from Bristol-Myers Squibb, Novartis, Sanofi, and Genentech. He has been a consultant for Bayer Pharmaceuticals, Repertoire Inc, Bristol Myers Squibb, Compass Therapeutics, EMD Serono, Genentech, Novartis Pharmaceuticals, and Sanofi Genzyme.

## ASUPPLEMENTAL INFORMATION TITLES AND LEGENDS

1. Supplemental Figures 1-10 and Legends
2. Experimental Model, Study Participant Details and Method Details

## AUTHOR CONTRIBUTIONS

Conceptualization: IMPACC Network, PMB, SK, LIRE, LG

Data Curation: AH, CS, BP, JD-A

Formal analysis: GG, JM, JPG, JFM, AH, CS, TC, NDJ

Funding Acquisition: IMPACC Network

Methodology: GG, JM, JPG, JFM, SHK, LG

Software: GG, JM, JPG, JFM, SHK, LG

Resources: IMPACC Network

Supervision: SF, MCA, OL, KKS, THH, RRM, JD-A, SHK, LG, LIRE

All authors wrote, edited, and reviewed the manuscript.

## Notes

https://bitbucket.org/kleinstein/impacc-public-code/src/master/multiomics-longcovid/

