## Supplementary material for "Identification of a multi-omics factor predictive of long COVID in the IMPACC study": Gabernet Supplemental Figures

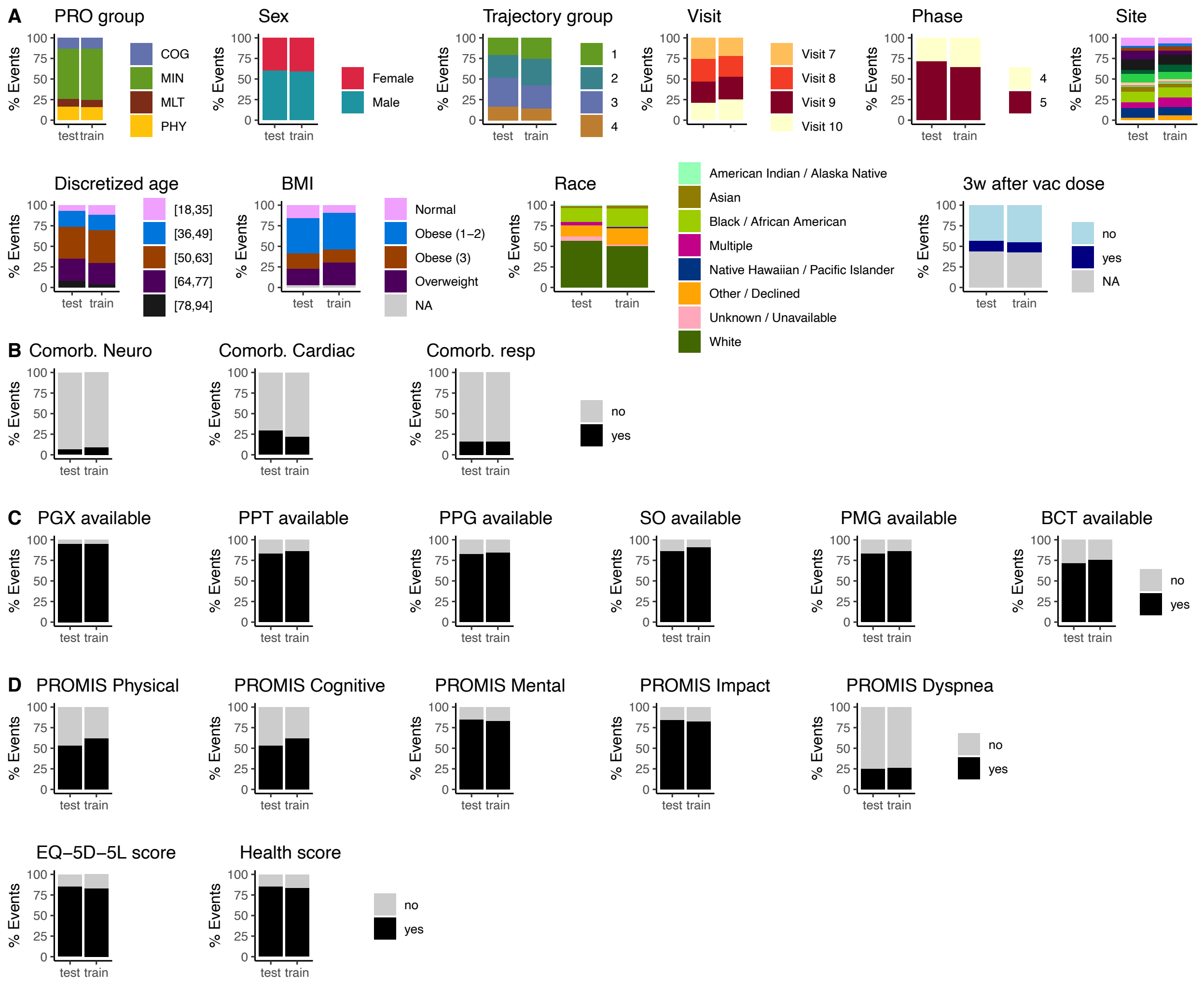


**Figure S1. IMPACC convalescent train and test cohort split evaluation. (A)** Balance of clinical characteristics within the Train and Test cohorts events including PRO group, sex (physician determined or reported sex at birth), trajectory group (acute COVID-19 disease severity measure), visit (target point of sample collection), phase (sample collection batch), enrollment site, discretized admit age, body mass index (BMI) and whether the sample was collected 3 weeks after a reported vaccine dose (3w after vac dose). **(B)** Balance of comorbidities (neurological, cardiac and respiratory) among participants in the train and test cohorts. **(C)** Availability of immunophenotyping assay data among participants in the train and test cohorts. **(D)** Availability of PROMIS scale scores among participants in the train and test cohorts.

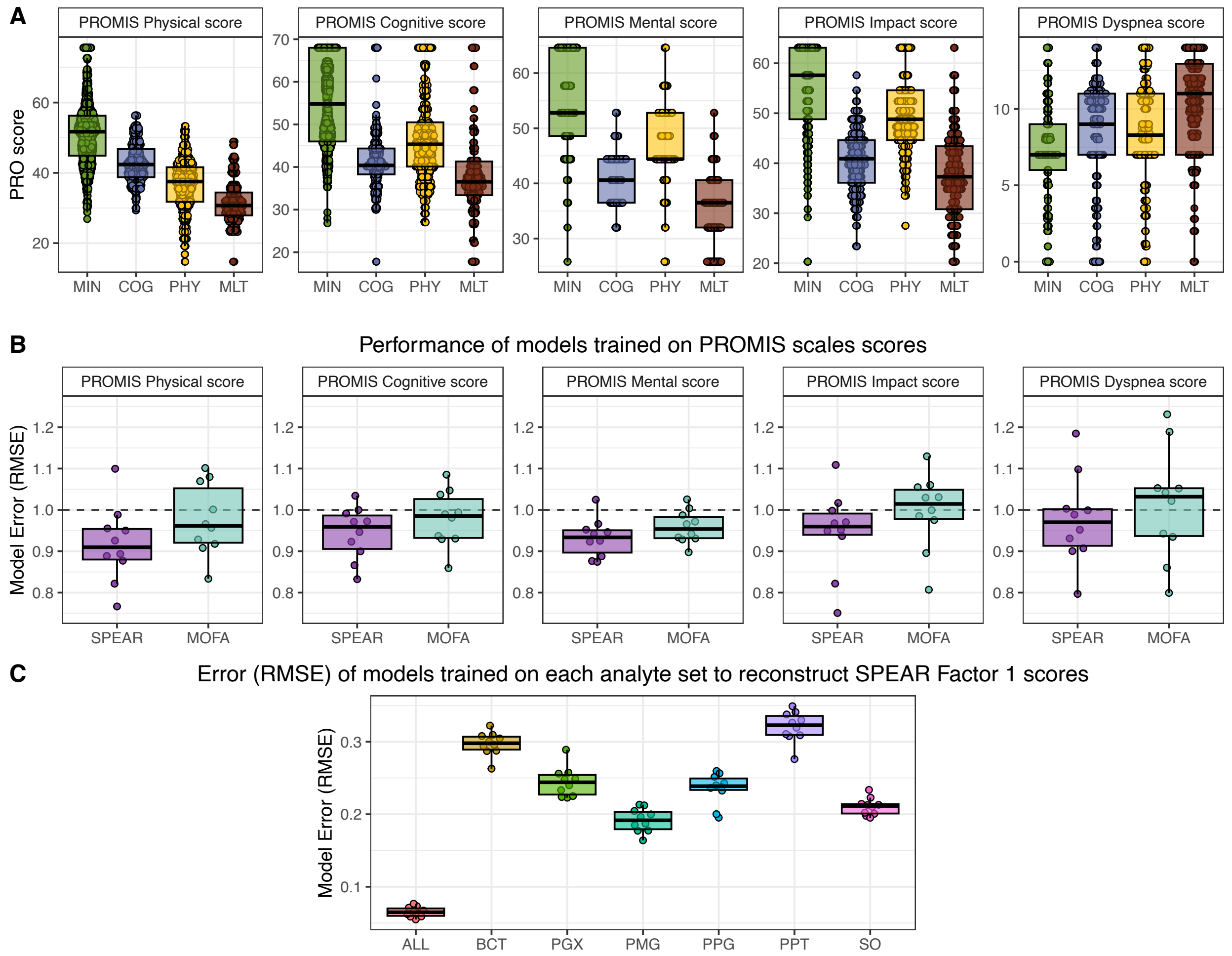


**Figure S2. Evaluation of SPEAR and MOFA models trained on PROMIS scale scores. (A)** PROMIS scale scores values for each PRO group. **(B)** Model error evaluated as root mean squared error (RMSE) of lasso regression models trained on SPEAR and MOFA factors using 10-fold cross-validation on the train cohort with each of the PROMIS scale scores as response variable (lower RMSE values indicate a better model). **(C)** Model error (RMSE) of lasso regression models trained on each individual analyte layer and all analytes (ALL) to reconstruct SPEAR Factor 1 scores. Individual analyte layers include Blood CyTOF MSI values (BCT), PBMC transcriptomics (PGX), plasma metabolomics global (PMG), plasma proteomics global (PPG), plasma proteomics targeted (PPT) and serum O-link (SO).

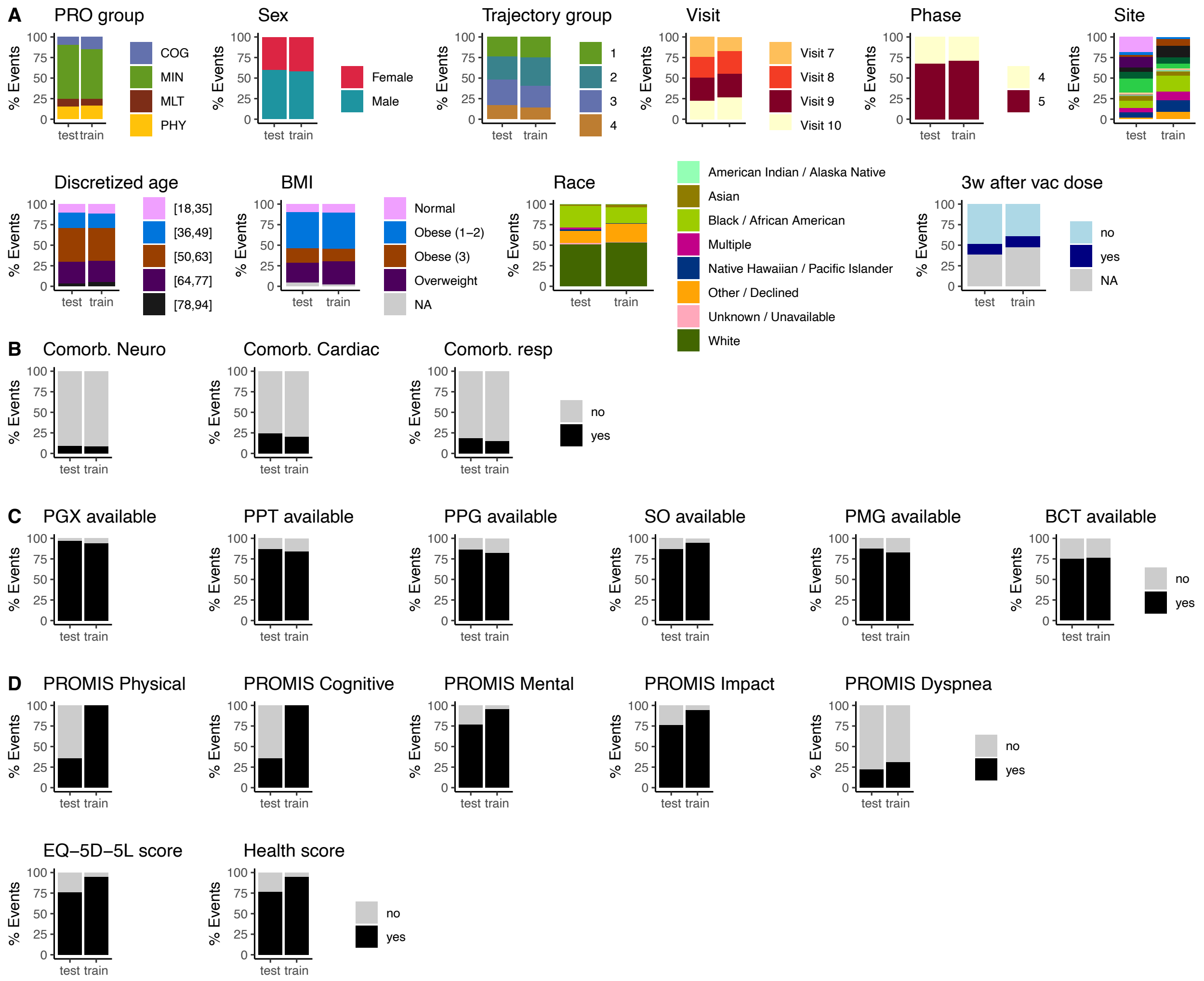


**Figure S3. IMPACC convalescent train and extended test set split evaluation.** To associate the recovery factor with LC vs MIN participants or PRO groups we considered an extended test set which comprised all participants in the Test cohort (with and without PROMIS Physical score) and participants in the Train cohort which did not have a PROMIS Physical score available for any of their measured samples and thus were not included in model training. The balance of the clinical characteristics **(A)**, comorbidities **(B)**, immunophenotyping assay data availability **(C)** and PRO measure score availability **(D)** of these two sets is shown.

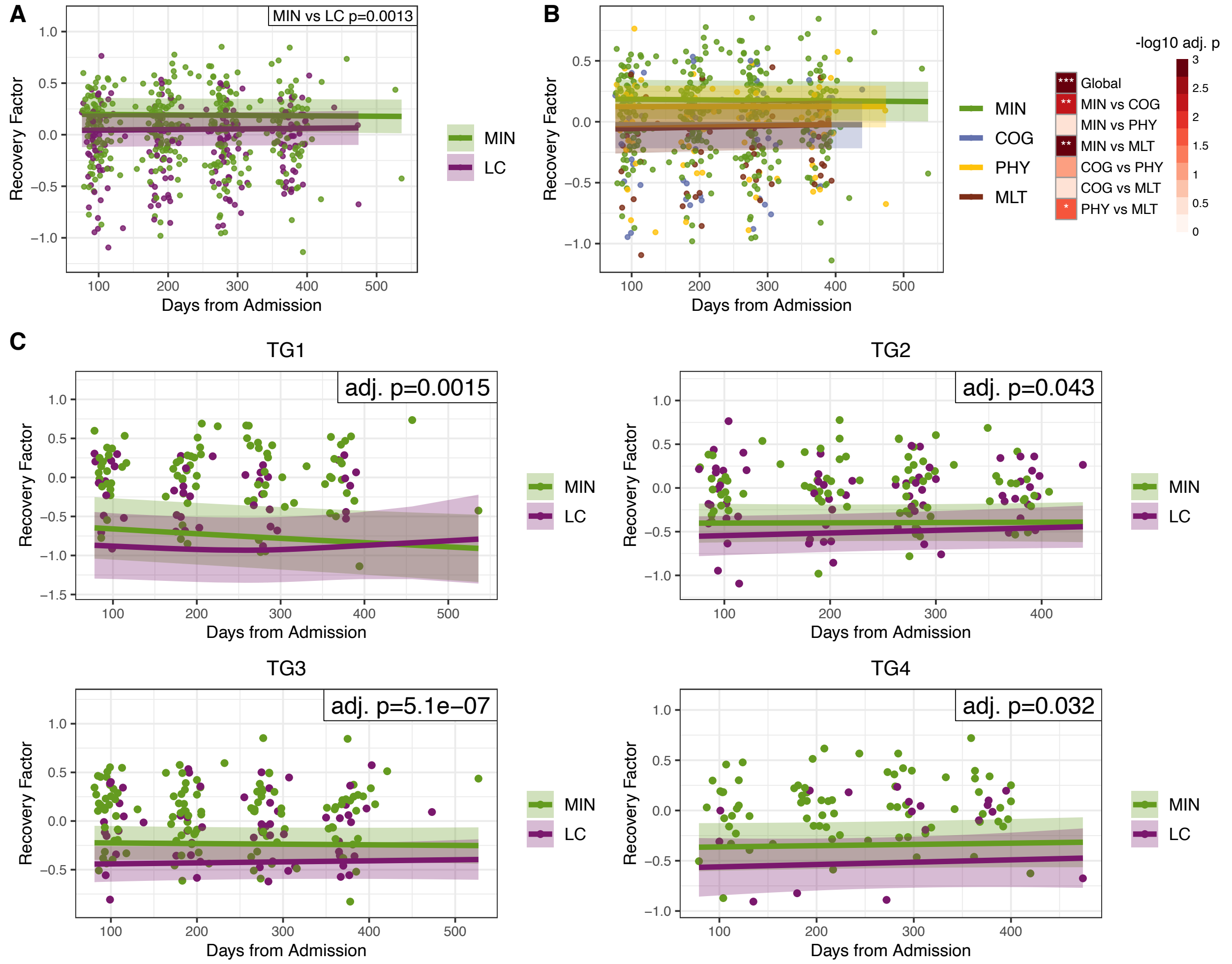


**Figure S4. Recovery factor is longitudinally associated with long COVID. (A)** Recovery factor scores for extended Test set MIN and LC participants. The p-value shows the longitudinal association of the recovery factor scores with MIN vs LC. **(B)** Recovery factor scores for the test cohort participants for each PRO group (MIN, COG, PHY, MLT). The heatmap shows the longitudinal association of the recovery factor with the PRO groups when considering global differences across all groups (Global) and pairwise differences among PRO groups. **(C)** Recovery factor scores and their association with MIN vs LC, subset by acute trajectory group.

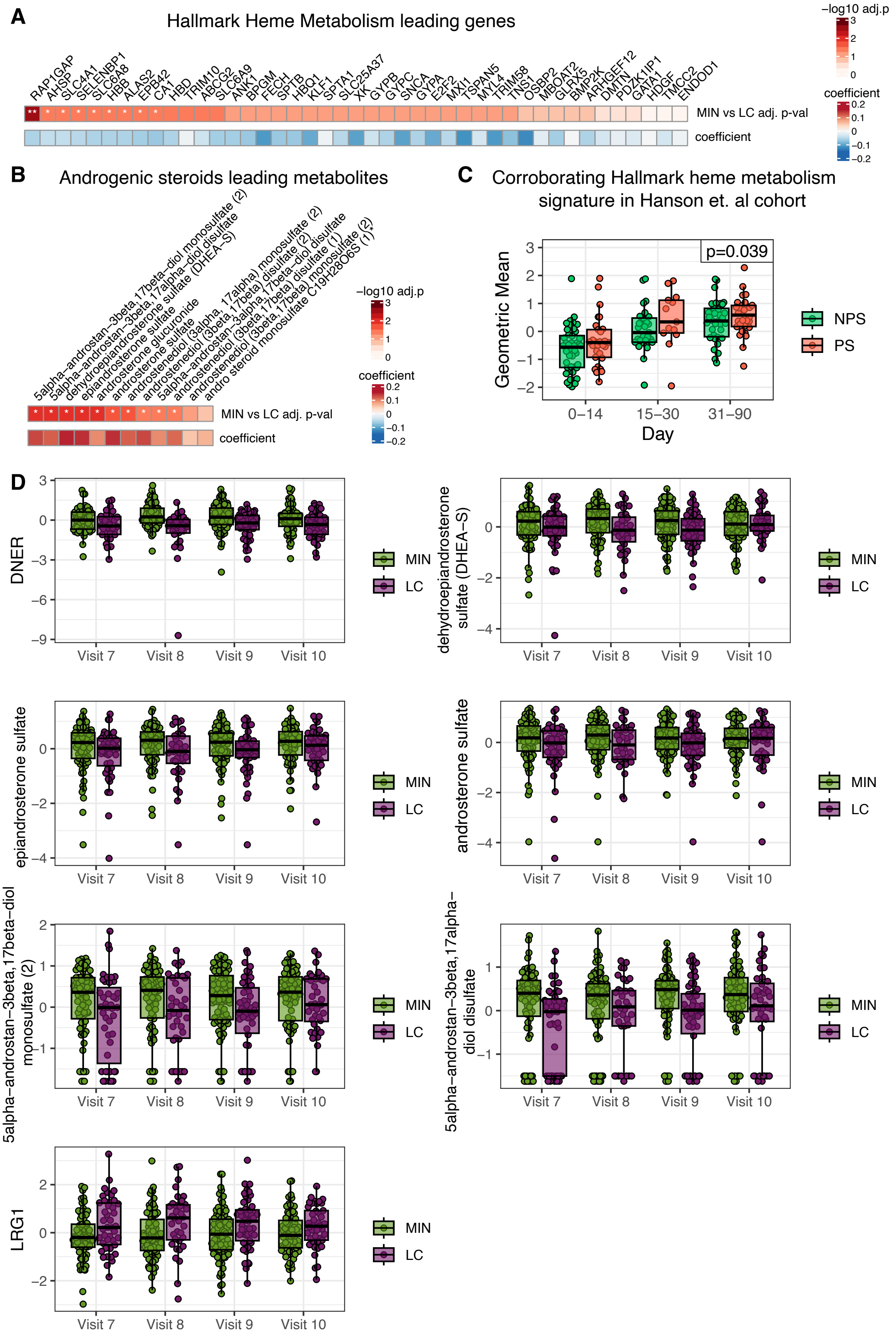
A
Hallmark Heme Metabolism leading genes
RAP1GAP AHSP SLC4A1 SELENBP1 SLC6A8 HBB ALAS2 EPB42 CA1 HBD TRIM10 ABCG2 SLC6A9 ANK1 BPGM FECH SPTB HBQ1 KLF1 SPTA1 SLC25A37 XK GYPB GYPC SNCA GYPA E2F2 MXI1 TSPAN5 MYL4 TRIM58 TNS1 OSBP2 MBOAT2 GLRX5 BMP2K ARHGEF12 DMTN PDZK1IP1 GATA1 HDGF TMCC2 ENDOD1
MIN vs LC adj. p-val
coefficient
−log10 adj.p
B
Androgenic steroids leading metabolites
5alpha-androstan-3beta,17beta-diol monosulfate (2)
5alpha-androstan-3beta,17alpha-diol disulfate
dehydroepiandrosterone sulfate (DHEA-S)
epiandrosterone sulfate
androsterone glucuronide
androsterone sulfate
androstenediol (3alpha, 17alpha) monosulfate (2)
androstenediol (3beta,17beta) disulfate (2)
5alpha-androstan-3alpha,17beta-diol disulfate
androstenediol (3beta,17beta) disulfate (1)
androstenediol (3beta,17beta) monosulfate (2)
andro steroid monosulfate C19H28O6S (1)
MIN vs LC adj. p-val
coefficient
C
Corroborating Hallmark heme metabolism signature in Hanson et. al cohort
p=0.039
Geometric Mean
Day
0–14
15–30
31–90
NPS
PS
D
DNER
dehydroepiandrosterone sulfate (DHEA−S)
epiandrosterone sulfate
androsterone sulfate
5alpha−androstan−3beta,17beta−diol monosulfate (2)
5alpha−androstan−3beta,17alpha−diol disulfate
LRG1
Visit 7
Visit 8
Visit 9
Visit 10
MIN
LC

**Figure S5. Individual associations of analytes from pathways associated with the recovery factor or significant Spear analytes with MIN vs LC status, and evaluation of the recovery factor Hallmark heme metabolism signature in an external cohort. (A)** Longitudinal association of the Hallmark heme metabolism pathway set leading genes with MIN vs LC (top; adj. intercept p-value), and SPEAR factor loadings (bottom; coefficient in the factor). **(B)** Longitudinal association of the Androgenic steroids set leading metabolites with MIN vs LC (top; adj. intercept p-value), and SPEAR factor loadings (bottom; coefficient in the factor). For panels A-B, (* adj. p-value < 0.05, ** adj. p-value < 0.01). (**C**) The Hallmark heme metabolism geometric mean signature was evaluated in whole blood transcriptomics measurements from participants with persisting symptoms (PS) and no persisting symptoms (NPS) in the Hanson et al. (2024)[22] cohort. (**D**) Significant SPEAR analyte values for the Test cohort in the MIN vs LC groups at the convalescent phase visits. Plots are shown only for Significant SPEAR analytes that were also significantly associated with MIN vs LC status (Figure 3B).

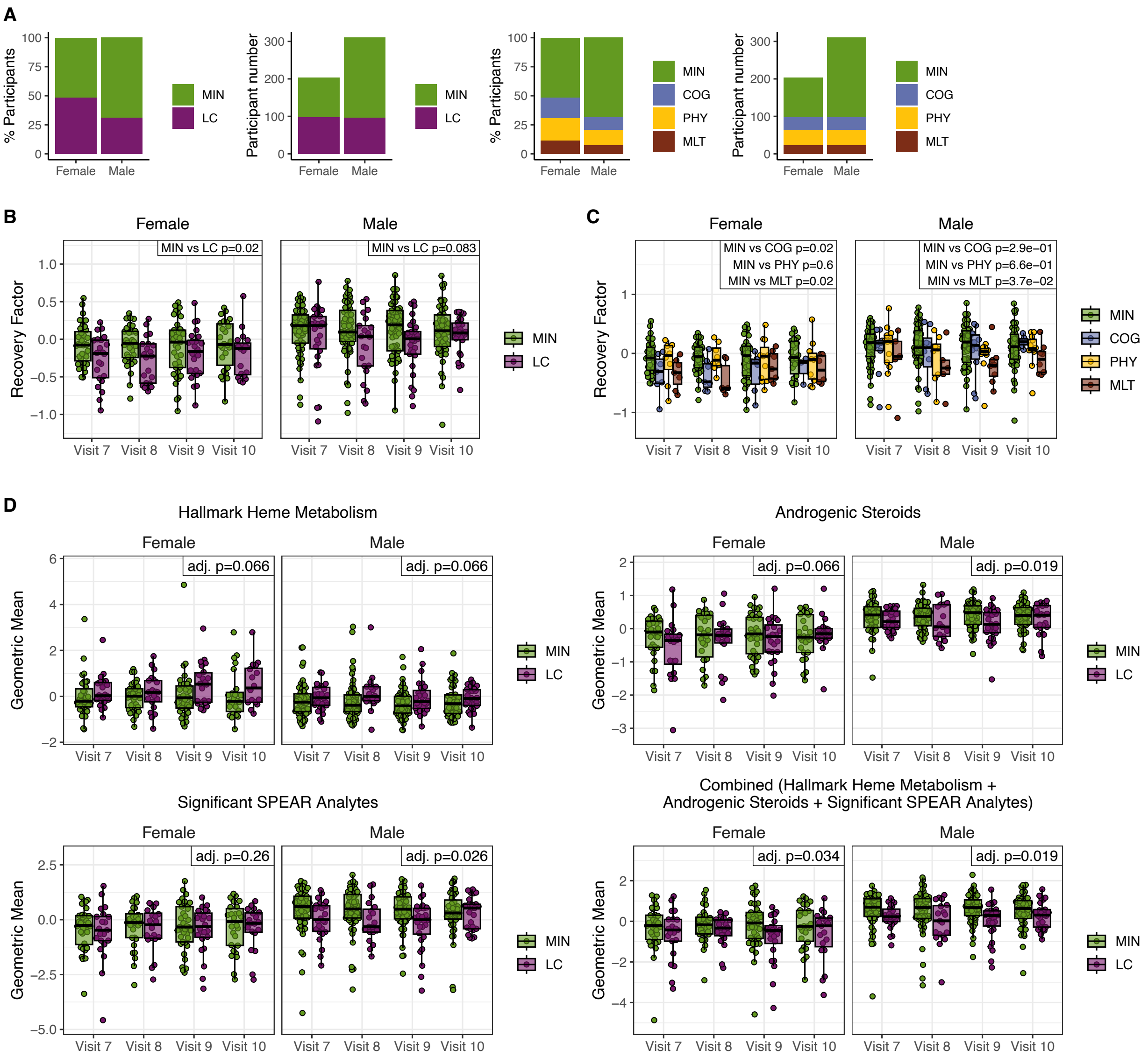


**Figure S6. Effects of sex on the recovery factor. (A)** Percentage and number participants in the MIN and LC group as well as individual PRO groups for Females and Males in the Convalescent Cohort. **(B)** Recovery factor scores for the Test cohort for MIN and LC groups stratified by sex. **(C)** Recovery factor scores for the Test cohort for PRO groups stratified by sex. **(D)** Test cohort geometric mean scores stratified by sex. Adjusted p-values show the association of the recovery scores with MIN vs LC. Associations were performed separately for males and females for each score.

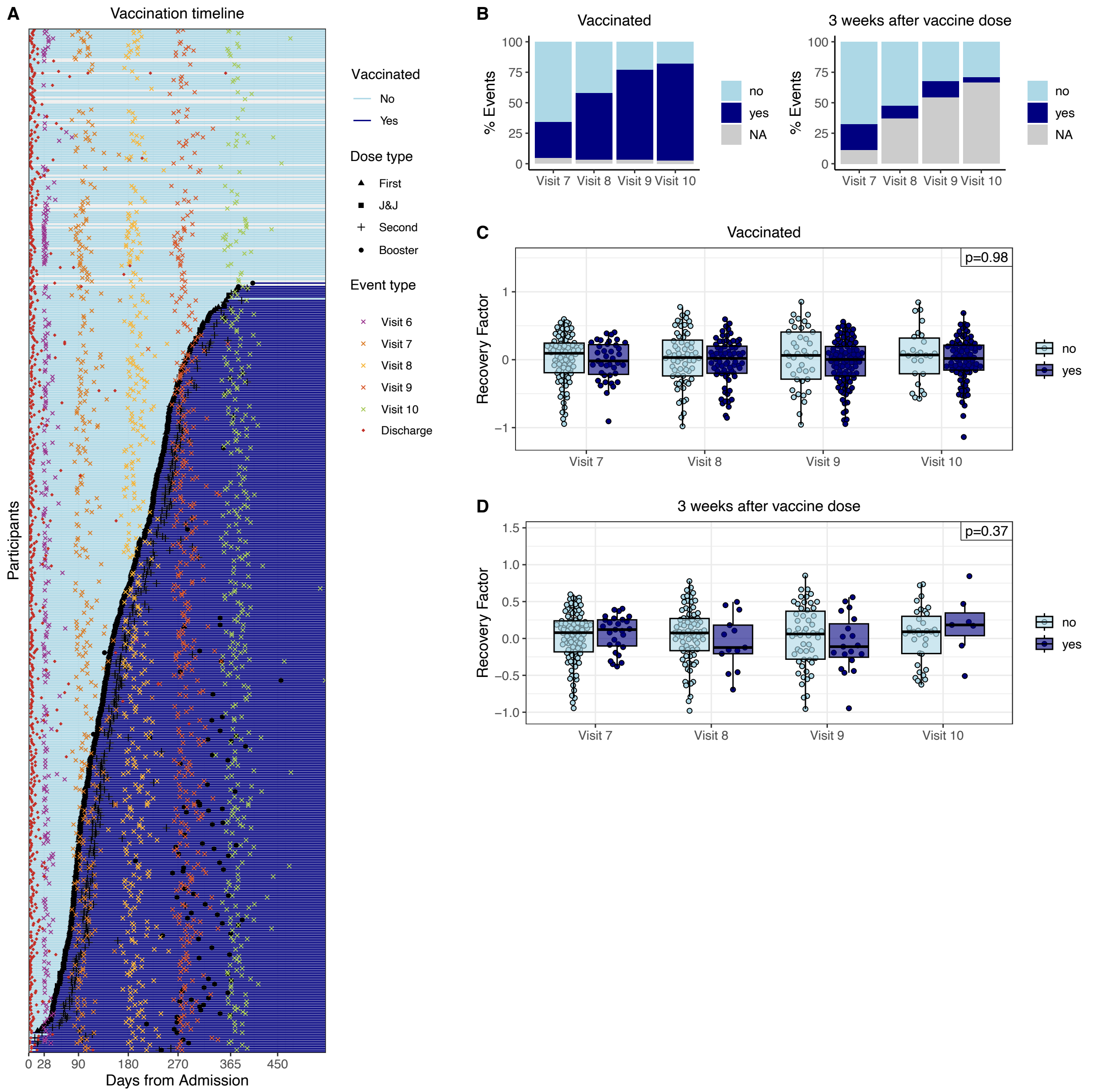


**Figure S7. Effects of vaccination on the recovery factor. (A)** Outpatient visits, hospital discharge and SARS-CoV-2 vaccination events timeline in days from admission for each participant in the Convalescent Cohort. Participants were ordered according to their first vaccination dose. **(B)** Percentage of events in each visit occurring after the participant received the first vaccine dose (vaccinated) or within 3 weeks after any vaccine dose. NA indicate events where the participant vaccination status, or exact vaccination date is unknown. **(C)** Recovery factor scores for events occurring before and after a participant received their first vaccine dose. **(D)** Recovery factor scores for events occurring within 3 weeks after any vaccine dose. P-values indicate the association of the recovery factor scores with the vaccination status.

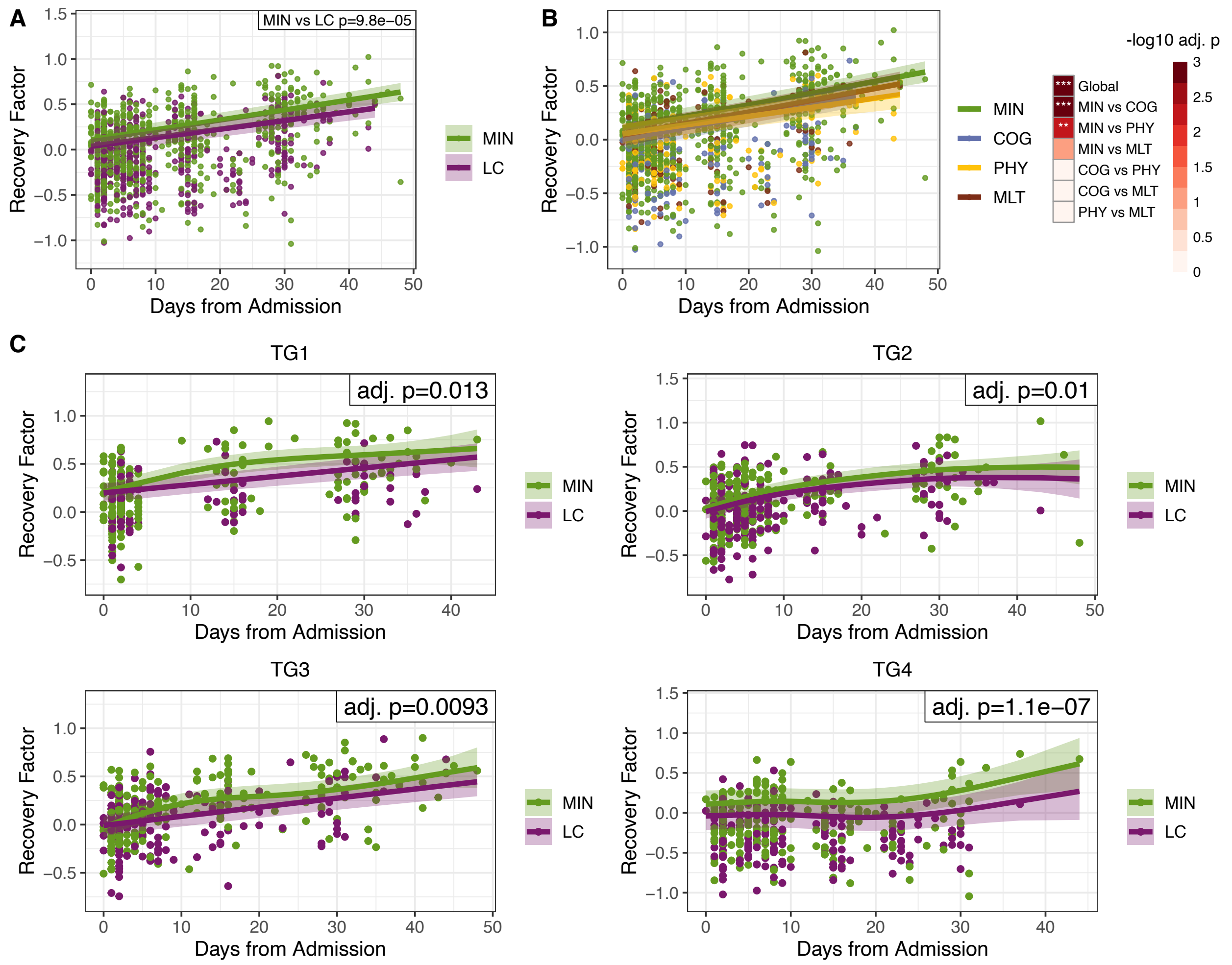


**Figure S8. Recovery factor is longitudinally associated with long COVID already during the acute phase. (A)** Recovery factor scores for MIN and LC participants during the acute disease phase. **(B)** Recovery factor scores for PRO group participants during the acute disease phase. The heatmap shows the longitudinal association of the recovery factor with the PRO groups when considering global differences across all groups (ALL PRO) and pairwise differences among PRO groups (*** adj. p < 0.001, ** adj. p < 0.01). **(C)** Recovery factor scores during the acute phase and their association with MIN vs LC, subset by acute trajectory group. Adjusted p-values indicate the association of the recovery factor scores with MIN vs LC.

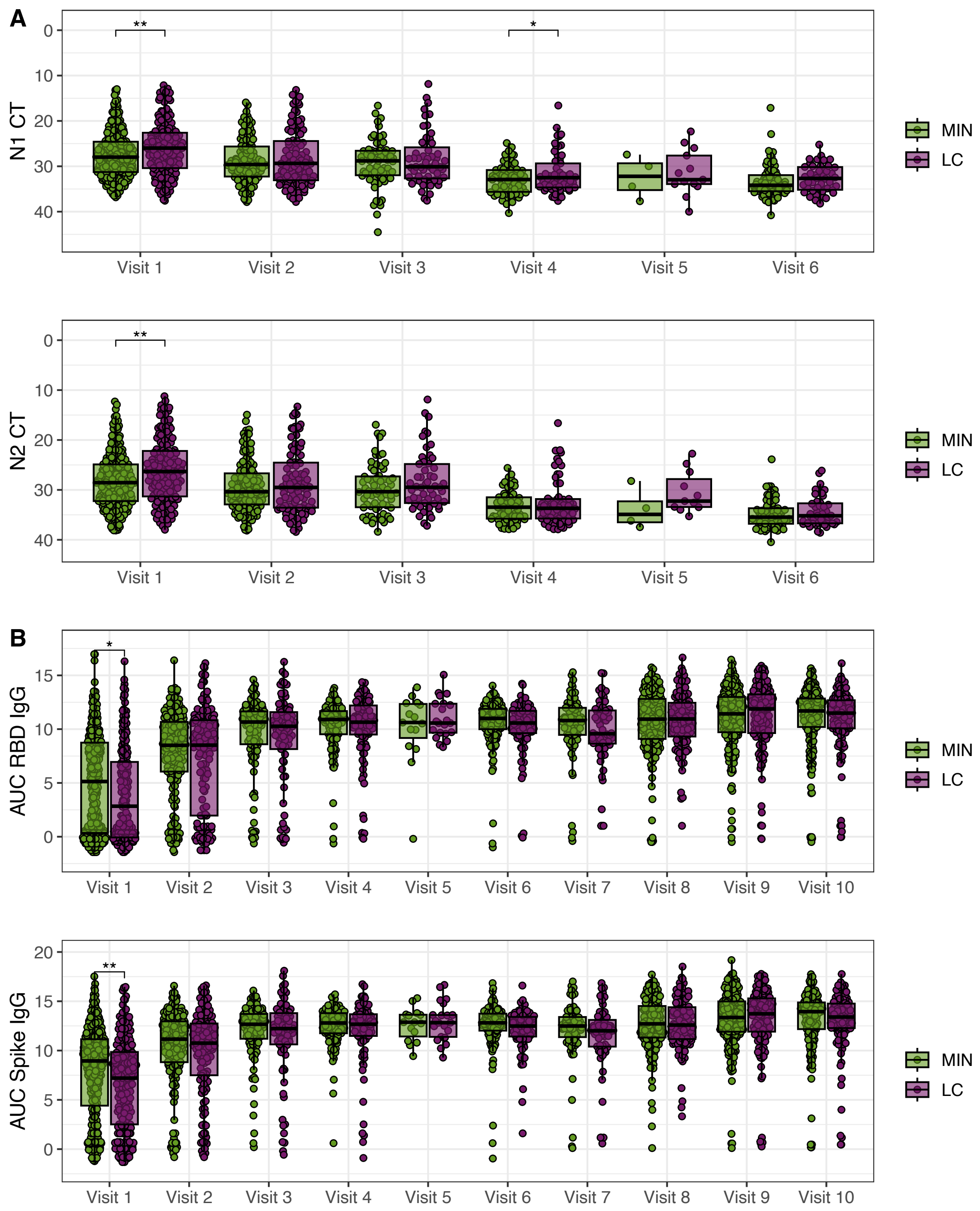


**Figure S9. Viral load and antibody titers at Visit 1 associate with LC status in the convalescent period. (A)** SARS-CoV-2 N1 and N2 gene PCR cycle threshold (Ct) values indicating viral load for the MIN and LC participant groups. Lower Ct values indicate higher viral loads, so the y axis is reversed. **(B)** Antibody titers against the SARS-CoV-2 virus full Spike protein (Spike IgG) and Receptor Binding Domain (RBD IgG). Area under the curve (AUC) values for the MIN and LC participant groups are shown.

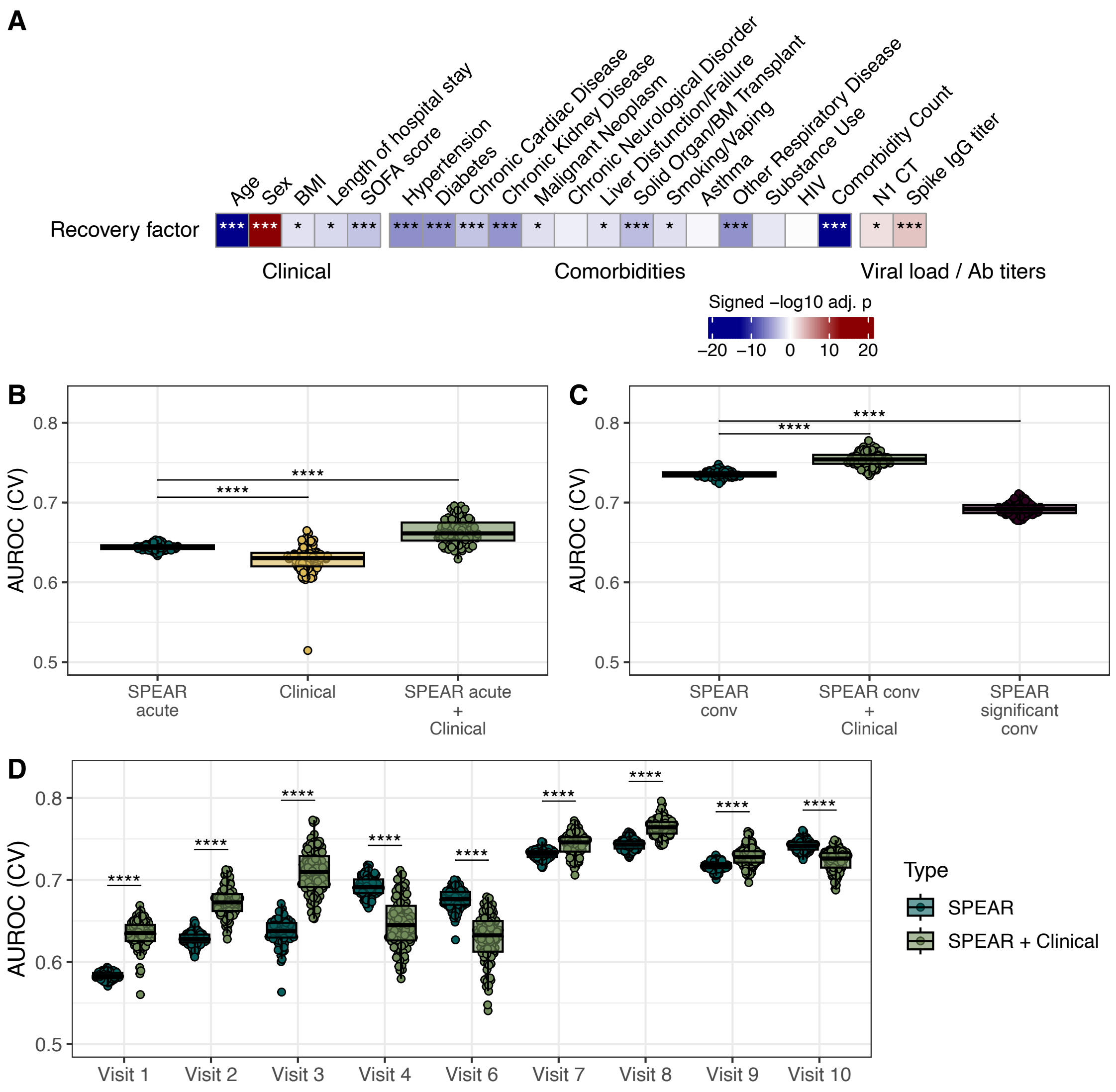
A
Age
Sex
BMI
Length of hospital stay
SOFA score
Hypertension
Diabetes
Chronic Cardiac Disease
Chronic Kidney Disease
Malignant Neoplasm
Chronic Neurological Disorder
Liver Disfunction/Failure
Solid Organ/BM Transplant
Smoking/Vaping
Asthma
Other Respiratory Disease
Substance Use
HIV
Comorbidity Count
N1 CT
Spike IgG titer
Recovery factor
Clinical
Comorbidities
Viral load / Ab titers
Signed −log10 adj. p
−20 −10 0 10 20
B
AUROC (CV)
SPEAR acute
Clinical
SPEAR acute + Clinical
C
SPEAR conv
SPEAR conv + Clinical
SPEAR significant conv
D
Visit 1
Visit 2
Visit 3
Visit 4
Visit 6
Visit 7
Visit 8
Visit 9
Visit 10
Type
SPEAR
SPEAR + Clinical

**Figure S10. Machine learning models based on the recovery factor scores at the acute and the convalescent phase together with clinical features predict LC status during the convalescent phase. (A)** Heatmap of signed adjusted p-values indicating the clinical feature term significance from a linear mixed-effect model with enrollment site and participant as random effects to explain the convalescent phase recovery factor scores. Sex and discretized age were further adjusted as fixed effects for clinical features other than sex and age. Color represents the term directionality. **(B)** Predictive performance of lasso classification models trained on the average of the recovery factor scores in the acute phase (SPEAR acute), the clinical variables shown in A (Clinical), as well as the combination of the recovery factor scores in the acute phase and clinical features (SPEAR acute + Clinical). **(C)** Predictive performance of lasso classification models trained on the average of the recovery factor scores in the convalescent phase (SPEAR conv), the combination of the recovery factor scores in the convalescent phase and the clinical variables shown in A (Spear conv + Clinical), and a model trained on the average values over the convalescent visits of the 26 SPEAR significant analytes (SPEAR significant conv). **(D)** Predictive performance of lasso models trained on the recovery factor scores at each of the acute (1-6) and convalescent (7-10) visits, compared to the same models including the clinical features in A. Visit 5 was omitted as there were insufficient measurements for the prediction evaluation. For panels B-D the mean AUROC of a 10-fold cross-validation on the Train Cohort, for 100 bootstrapped model training repetitions are shown. Significance was calculated by standard normal approximation of bootstrapped differences between models (t-test, ****adj. p-value ≤ 0.0001).
