## Supplementary material for "Identification of a multi-omics factor predictive of long COVID in the IMPACC study": Gabernet Supplemental Methods

### EXPERIMENTAL MODEL AND STUDY PARTICIPANT DETAILS

#### Ethics

NIAID staff conferred with the Department of Health and Human Services Office for Human Research Protections (OHRP) regarding the potential applicability of the public health surveillance exception [45CFR46.102(l) (2)] to the IMPACC study protocol. OHRP concurred that the study satisfied criteria for the public health surveillance exception, and the IMPACC study team sent the study protocol, and participant information sheet for review and assessment to institutional review boards (IRBs) at participating institutions. Twelve institutions elected to conduct the study as public health surveillance, while 3 sites with prior IRB-approved biobanking protocols elected to integrate and conduct IMPACC under their institutional protocols (University of Texas at Austin, IRB 2020-04-0117; University of California San Francisco, IRB 20-30497; Case Western Reserve University, IRB STUDY20200573) with informed consent requirements. Participants enrolled under the public health surveillance exclusion were provided information sheets describing the study, samples to be collected, and plans for data de-identification and use. Those who requested not to participate after reviewing the information sheet were not enrolled. In addition, participants did not receive compensation for study participation while inpatient, and subsequently were offered compensation during outpatient follow-ups.

#### IMPACC Cohort characteristics

The IMPACC Cohort consists of 1164 participants admitted to 20 US hospitals (affiliated with 15 academic institutions) between May 2020 and March 2021 that were enrolled within 72h of hospital admission for COVID-19 infection. Participants with confirmed positive SARS-CoV-2 PCR and COVID-19 infection symptomatology were followed longitudinally during the acute infection phase (1-28 days after hospital admission) and the convalescent phase (3 months to 12 months after hospital admission). Details on the study design for clinical data and biological sample collection were previously described[1–3]. Clinical data including length of hospital stay, complications, mortality, and other pre-defined outcomes were collected over the acute phase. Self-reported symptoms, reinfections, SARS-CoV-2 vaccination, rehospitalizations, and standardized patient-reported outcome surveys were assessed quarterly over the convalescent period through a mobile application and remote visits[1]. The surveys at these remote visits captured upper respiratory symptoms, cardiopulmonary symptoms, systemic symptoms, neurologic symptoms, and gastrointestinal symptoms. Additionally, six validated Patient-Reported Outcome (PRO) measures were used to evaluate general health and deficits in specific health domains including EQ-5D-5L[4], and the Patient-Reported Outcomes Measurement Information System (PROMIS)[5,6] Physical function, Cognitive function, Global Health Mental, Psychosocial Illness Impact and Dyspnea Time Extension surveys. Overall health was also assessed by a health recovery score to indicate overall post discharge mental and physical function compared to pre-COVID function[1].

**PRO group assignments**

Participants were assigned to PRO groups as previously described[1]. Briefly, all six PRO scores were standardized using standard PROMIS T-scores or a standard value set. The health recovery score was computed utilizing a Visual Analog Scale ranging from 1 to 100. The individual PRO scores collected over the convalescent period were modelled longitudinally using Latent Class Mixed Models (LCMM). A Ward clustering analysis was then applied to group participants with similar PRO longitudinal patterns. Four distinct clusters were identified, and t-statistics were calculated comparing the mean value of individual PRO scores within each cluster versus the mean value of that PRO across the remaining clusters. After association with the specific PROs, clusters were labeled minimal deficit (MIN), physical predominant (PHY), mental/cognitive predominant (COG), and multi/pan domain deficit (MLT).

**IMPACC Convalescent cohort definition**

In this study, we only included participants with at least one quarterly set of surveys post discharge and immune profiling measurements for at least one of the following assays during the convalescent phase (visits 7-10 at 3, 6, 9, and 12 months post hospital discharge): serum O-link, plasma targeted proteomics, plasma global proteomics, plasma global metabolomics, PBMC transcriptomics and CyTOF measurements, yielding an analysis cohort of 513 participants (Table S1).

**Table S1**. Demographics and baseline characteristics of the Convalescent cohort

| | | **Overall** (N=513) | **MIN** (n=318) | **LC** (n=195) | **Overall p-value** |
|---|---|---|---|---|---|
| Age at enrollment (years), median (IQR) | (n=513) | 57.0 (19.0) | 56.5 (20.0) | 57.0 (16.0) | 0.653 |
| Sex at birth | Male | 310 (60%) | 213 (67%) | 97 (50%) | <.001 |
| | Female | 203 (40%) | 105 (33%) | 98 (50%) | |
| Race | White | 248 (48%) | 157 (49%) | 91 (47%) | 0.352 |
| | Black | 118 (23%) | 73 (23%) | 45 (23%) | |
| | Other | 101 (20%) | 56 (18%) | 45 (23%) | |
| | Asian | 19 (4%) | 13 (4%) | 6 (3%) | |
| | Multiple | 7 (1%) | 6 (2%) | 1 (1%) | |
| | American Indian/Alaska Native | 6 (1%) | 2 (1%) | 4 (2%) | |
| | Native Hawaiian/Pacific Islander | 4 (1%) | 3 (1%) | 1 (1%) | |
| | Unknown | 10 (2%) | 8 (3%) | 2 (1%) | |
| Hispanic ethnicity | Non-Hispanic | 334 (65%) | 208 (65%) | 126 (65%) | 0.717 |
| | Hispanic | 168 (33%) | 102 (32%) | 66 (34%) | |
| | Unknown | 11 (2%) | 8 (3%) | 3 (2%) | |
| Comorbidities | Hypertension | 282 (55%) | 163 (51%) | 119 (61%) | 0.031 |
| | Diabetes | 168 (33%) | 94 (30%) | 74 (38%) | 0.049 |
| | Chronic respiratory (not asthma) | 85 (17%) | 37 (12%) | 48 (25%) | <.001 |
| | Asthma | 90 (18%) | 51 (16%) | 39 (20%) | 0.252 |
| | Chronic cardiac disease | 122 (24%) | 62 (19%) | 60 (31%) | 0.004 |
| | Chronic kidney disease | 62 (12%) | 42 (13%) | 20 (10%) | 0.32 |
| | Malignant neoplasm | 37 (7%) | 26 (8%) | 11 (6%) | 0.281 |
| | Chronic neurologic disorder | 49 (10%) | 20 (6%) | 29 (15%) | 0.001 |
| | Liver disease | 25 (5%) | 12 (4%) | 13 (7%) | 0.14 |
| | History of SOT or BMT | 38 (7%) | 18 (6%) | 20 (10%) | 0.054 |
| | Current or former smoking and/or vaping | 151 (29%) | 86 (27%) | 65 (33%) | 0.129 |
| | Substance use (drugs, alcohol, and/or cannabis) | 35 (7%) | 20 (6%) | 15 (8%) | 0.541 |
| BMI Category | Underweight | 4 (1%) | 4 (1%) | 0 (0%) | 0.003 |
| | Normal weight | 54 (11%) | 34 (11%) | 20 (10%) | |
| | Overweight (25.1-29.9) | 141 (27%) | 95 (30%) | 46 (24%) | |
| | Class 1-2 Obesity (30-39.9) | 220 (43%) | 135 (42%) | 85 (44%) | |
| | Class 3 Obesity (40+) | 80 (16%) | 37 (12%) | 43 (22%) | |
| | Missing | 14 (3%) | 13 (4%) | 1 (1%) | |
| Number of comorbidities | 0 | 34 (7%) | 29 (9%) | 5 (3%) | <.001 |
| | 1 | 82 (16%) | 62 (19%) | 20 (10%) | |
| | 2 | 100 (19%) | 66 (21%) | 34 (17%) | |
| | 3 | 101 (20%) | 55 (17%) | 46 (24%) | |
| | 4 | 69 (13%) | 39 (12%) | 30 (15%) | |
| | 5 | 127 (25%) | 67 (21%) | 60 (31%) | |
| Infiltrates on chest X-ray or chest tomography | No infiltrates | 110 (22%) | 67 (22%) | 43 (23%) | 0.978 |
| | Unilateral infiltrates | 45 (9%) | 29 (10%) | 16 (9%) | |
| | Bilateral infiltrates | 331 (68%) | 206 (68%) | 125 (68%) | |
| | Unknown/Missing | 3 (1%) | 2 (1%) | 1 (1%) | |
| Level of respiratory support | Mechanically ventilated, or ECMO (OS=6) | 33 (6%) | 20 (6%) | 13 (7%) | 0.022 |
| | Non-invasive ventilation, or high flow nasal O2 (OS=5) | 77 (15%) | 58 (18%) | 19 (10%) | |
| | Supplemental oxygen (not high flow) (OS=4) | 290 (57%) | 180 (57%) | 110 (56%) | |
| | None (OS=3) | 113 (22%) | 60 (19%) | 53 (27%) | |
| SpO2/FiO2 ratio category at lowest sat | 235 or lower | 84 (16%) | 62 (19%) | 22 (11%) | 0.077 |
| | 236-315 | 96 (19%) | 57 (18%) | 39 (20%) | |
| | 315 or higher | 304 (59%) | 184 (58%) | 120 (62%) | |
| | Missing | 29 (6%) | 15 (5%) | 14 (7%) | |
| SOFA Score, median (IQR) | (n=513) | 0.0 (2.0) | 0.0 (2.0) | 0.0 (2.0) | 0.991 |
| Lymphocyte count (1000s/microliter), median | (n=431) | 1.0 (0.9) | 1.0 (0.8) | 1.2 (1.1) | 0.234 |
| Platelets (1000s/microliter), median (IQR) | (n=487) | 239.0 (124.0) | 239.0 (126.0) | 238.0 (125.5) | 0.023 |
| ALT (Units/L), median (IQR) | (n=453) | 32.0 (34.0) | 33.5 (34.0) | 32.0 (32.0) | 0.271 |
| Creatinine (mg/dL), median (IQR) | (n=494) | 0.9 (0.4) | 0.9 (0.4) | 0.8 (0.4) | 0.596 |
| CRP (mg/L), median (IQR) | (n=375) | 13.0 (61.1) | 13.2 (55.0) | 12.3 (64.1) | 0.23 |
| D-dimer (mg/L), median (IQR) | (n=369) | 0.7 (0.8) | 0.8 (0.9) | 0.6 (0.6) | 0.063 |
| Troponin (ng/mL), median (IQR) | (n=170) | 0.0 (0.1) | 0.0 (0.1) | 0.0 (0.1) | 0.899 |
| Length of stay (days), median (IQR) | (n=500) | 5.0 (5.0) | 5.0 (6.0) | 6.0 (5.0) | 0.206 |
| ICU at any time during acute hospitalization | Yes | 134 (26%) | 84 (26%) | 50 (26%) | 0.846 |
| Acute trajectory group | 1 | 126 (25%) | 83 (26%) | 43 (22%) | 0.254 |
| | 2 | 167 (33%) | 95 (30%) | 72 (37%) | |
| | 3 | 141 (27%) | 86 (27%) | 55 (28%) | |
| | 4 | 79 (15%) | 54 (17%) | 25 (13%) | |
| Complications | Any complications | 446 (87%) | 284 (89%) | 162 (83%) | 0.042 |
| | Number of complications, median (IQR) n=513 | 2.0 (2.0) | 2.0 (2.0) | 2.0 (2.0) | 0.367 |
| | Acute renal injury/ failure | 84 (16%) | 50 (16%) | 34 (17%) | 0.611 |
| | Liver dysfunction/ failure | 63 (12%) | 45 (14%) | 18 (9%) | 0.099 |
| | Anemia | 57 (11%) | 31 (10%) | 26 (13%) | 0.21 |
| | Shock (use of vasopressors) | 37 (7%) | 18 (6%) | 19 (10%) | 0.083 |
| | Bacteremia | 41 (8%) | 27 (8%) | 14 (7%) | 0.595 |
| | Atrial fibrillation | 25 (5%) | 15 (5%) | 10 (5%) | 0.834 |
| | Acute venous thromboembolism | 25 (5%) | 14 (4%) | 11 (6%) | 0.527 |
| | Congestive heart failure (CHF) / cardiomyopathy | 22 (4%) | 12 (4%) | 10 (5%) | 0.462 |
| | Hyperglycemia | 20 (4%) | 11 (3%) | 9 (5%) | 0.511 |
| Medications | Remdesivir | 332 (65%) | 211 (66%) | 121 (62%) | 0.322 |
| | Steroids | 354 (69%) | 229 (72%) | 125 (64%) | 0.06 |
| Convalescent symptoms | Any symptom reported during convalescence | 273 (53%) | 140 (44%) | 133 (68%) | <.001 |
| | Any upper respiratory | 91 (18%) | 34 (11%) | 57 (29%) | <.001 |
| | Conjunctivitis/red eyes | 65 (13%) | 23 (7%) | 42 (22%) | <.001 |
| | Sore throat | 39 (8%) | 13 (4%) | 26 (13%) | <.001 |
| | Any cardiopulmonary | 186 (36%) | 87 (27%) | 99 (51%) | <.001 |
| | Dyspnea | 155 (30%) | 69 (22%) | 86 (44%) | <.001 |
| | Cough | 106 (21%) | 43 (14%) | 63 (32%) | <.001 |
| | Any systemic | 154 (30%) | 61 (19%) | 93 (48%) | <.001 |
| | Fatigue | 96 (19%) | 31 (10%) | 65 (33%) | <.001 |
| | Myalgia | 106 (21%) | 36 (11%) | 70 (36%) | <.001 |
| | Fever | 14 (3%) | 6 (2%) | 8 (4%) | 0.135 |
| | Chills | 24 (5%) | 9 (3%) | 15 (8%) | 0.011 |
| | Any neurologic | 143 (28%) | 59 (19%) | 84 (43%) | <.001 |
| | Headache | 98 (19%) | 35 (11%) | 63 (32%) | <.001 |
| | Anosmia | 76 (15%) | 29 (9%) | 47 (24%) | <.001 |
| | Any gastrointestinal | 36 (7%) | 11 (3%) | 25 (13%) | <.001 |

### METHOD DETAILS

#### Sample processing and quantification and batch randomization

Biological sample collection and processing was homogeneously performed by every participating academic center. Batch randomization, sample processing and assay preparation and quantification were performed according to previously published protocols[2,3,7].

#### Train and test cohort split

Participants in the convalescent cohort were split into 80% Train cohort and 20% Test cohort, maintaining the proportions of PRO group individuals with each group.

#### Data preprocessing and additional quality control

Data preprocessing included sample filtering according to quality control steps, feature filtering according to remove features with a high percentage of missing values or low variance, batch correction, missing value imputation and data transformation (Table S2). We evaluated the influence of potential batch effects on the different assays using Principal variance component analysis (PVCA). The pre-processing steps for each data modality are described in detail below. Train and Test cohort datasets were independently processed to avoid data leakage between the datasets. For the feature filtering step, the same features were filtered on the Test cohort as identified on the Train cohort, to allow for the Factor score reconstruction.

**Table S2. Summary of data preparation steps.** For each assay, we first filtered out samples according to the sample filtering criteria and features based on the feature filtering criteria. N/A: no additional step taken. Pareto-scaling: in-house function of normalizing each centered variable by the square root of the standard deviation. Half-min: missing values were replaced using half the minimum of observed values for the corresponding feature.

| Assay name | Sample filtering | Feature filtering | Additional batch correction | Missing value imputation | Data transformation |
|---|---|---|---|---|---|
| PBMC gene expression (PGX) | Passed & questionable QC | Protein coding, Genes with CPM >=1 in >5% of samples in a PRO group, top 20% highly variable genes | RemoveBatchEffects (limma) | N/A | Log2CPM and scaling |
| Serum O-link (SO) | Passed & questionable QC | Remove samples with all missing values and features with >99% | N/A | N/A | Scaling |

| | | | | | |
|---|---|---|---|---|---|
| | | missing values | | | |
| Plasma proteomics global and targeted (PPG & PPT) | Passed & questionable QC | Removed features with >99% missing values | Batch correction (ComBAT) | Half-min | Median normalization in linear space Log2 transformation with pseudocount of 1 and scaling |
| Plasma metabolomics global (PMG) | Passed & questionable QC | Removed non-xenobiotic features with IQR=0 | | Half-min | Pareto-scaling |
| Blood CyTOF | Passed & questionable QC | Removed undefined features, debris, multiplets, platelets, RBC. Removed features with all missing values, and samples with >80% missing values. | N/A | N/A | Log1p transformation of blood cytof frequencies followed by scaling of cell subset across samples. |
| Blood CyTOF MSI (BCT) | Passed & questionable QC | Removed undefined features, debris, multiplets, platelets, RBC. Removed features with all missing values, and samples with more than 80% missing values. | RemoveBatchEffects (limma) | N/A | Scaling |

***PBMC Gene Expression (PGX)***

We filtered for protein-coding genes and removed genes with low expression (genes that did not exceed the threshold of counts per million (CPM) >= 1 in more than 5% of samples in at least one outcome group (PRO group). We also selected the top 20% highly variable features by mean absolute deviation (MAD) of the log-transformed CPM values. The count data was transformed using the *voom* function from the limma R package[8], batch correction was performed to remove the effects of technical variables, including sample processing batch (phase) and library preparation plate using the function *removeBatchEffect* from limma, and the data was scaled.

***Serum Olink (SO)***

Samples with all missing values and features with >99% missing values were removed, and the data was scaled.

***Plasma Proteomics Global and Targeted (PPG, PPT)***

Features with more than 99% missing values were removed, and the data was median normalized in lineal space, imputed using the half-min approach (missing values were replaced using half the minimum of observed values for the corresponding feature), log-transformed using a pseudocount of 1. Batch correction to correct for the effects of processing batches (phase) and plates was performed with the *ComBat* function[9] of the sva R package. Finally, data was scaled.

***Plasma Metabolomics Global (PMG)***

Missing values were imputed using the half-min approach, non-xenobiotic features with IQR=0 were removed as previously described[3], and the data was pareto-scaled.

***Whole Blood CyTOF cell frequencies (CyTOF)***

Cells were manually labeled into parent and child subsets as previously described[3] (Table S3). Cells annotated as cellular debris, multiplets, platelets, red blood cells, and undefined populations were removed. Cellular population counts were converted to a normalized frequency by dividing by the total counts per sample. For non-granulocyte cell types, the total counts normalization was calculated without granulocytes. The normalized frequencies were then log1p transformed and scaled by cell subset across samples.

**Table S3. Parent and child cell type annotations used in the whole blood CyTOF data.**

| Parent Cell Type | Child Cell Types |
|---|---|
| B Cell | B Cell (CD27+ non-switched memory), B Cell (CD27+ switched memory), B Cell (CD71+ activated), B Cell (Plasmablast), B Cell (naive), B Cell (transitional) |
| CD4+ T Cell | CD4+ NKT Cell, CD4+ T Cell (CM), CD4+ T Cell (EM CD27hi), CD4+ T Cell (EM CD27low), CD4+ T Cell (EM CD57hi), CD4+ T Cell (EMRA CD57hi), CD4+ T Cell (EMRA CD57low), CD4+ T Cell (activated), CD4+ T Cell (naive), CD4+ Treg (CD39hi), CD4+ Treg (CD39low), CD4+ Treg (naive) |
| CD8+ T Cell | CD8+ NKT Cell, CD8+ T Cell (CD161+ MAIT), CD8+ T Cell (CM), CD8+ T Cell (EM CD27hi), CD8+ T Cell (EM CD27low), CD8+ T Cell (EM CD57hi), CD8+ T Cell (EMRA CD57hi), CD8+ T Cell (EMRA CD57low), CD8+ T Cell (activated), CD8+ T Cell (naive) |
| γδ T Cell | γδ T Cell |
| NK Cell | NK Cell (CD56hi CD16low), NK Cell (CD56low CD16hi CD57hi), NK Cell (CD56low CD16hi CD57low) |
| Innate Lymphoid Cell | Innate Lymphoid Cell |
| Hematopoietic Progenitor Cell | Hematopoietic Progenitor Cell |

| | |
|---|---|
| Monocytes | Monocytes (CD14+CD16+), Monocytes (CD14+CD16-), Monocytes (CD14-CD16+) |
| Neutrophil | Neutrophil (CD16hi), Neutrophil (CD16low) |
| Eosinophil | Eosinophil |
| Basophil | Basophil |
| Conventional Dendritic Cell | Conventional Dendritic Cell |
| Plasmacytoid Dendritic Cell | Plasmacytoid Dendritic Cell |

#### *Whole Blood CyTOF marker MSI (BCT)*

Cells were filtered as described in the section above, and additionally DNA markers and bead size markers were removed. Features consisted of cell membrane protein markers mean signal intensity (MSI) per cell population. Features with more than 80% missing values were removed, and counts were scaled. Batch correction to correct for the effects of processing batches (phase) and plates was performed with the *removeBatchEffects* function from limma[8].

### Clinical data processing

#### *Convalescent symptom groups and longitudinal associations*

Symptoms reported at hospital admission (baseline) and during the convalescent period were categorized into three mutually exclusive groups: Cardiopulmonary (dyspnea, cough), systemic (fatigue, myalgia, chills, fever), and other (GI – nausea, vomiting; neurologic – anosmia, headache; upper respiratory – sore throat, red eyes/conjunctivitis). Reported symptoms within each group were counted to identify participants who had reported one of these symptoms during at least one convalescent timepoint, as well as participants who had reported a symptom at two or more timepoints, including baseline.

#### *Acute infection trajectory group definition*

Respiratory ordinal scores were assigned to each acute participant visit based on recorded clinical data, such as respiratory supports and hospitalization status[10]. Unsupervised clustering of these scores over time was used to group participants into five groups of disease severity[10]. The clustering was run twice: once for the initial clinical data analysis and again once all data collection and cleaning was complete, generating two sets of groups. This analysis uses the latter groups, based on the more complete dataset. These groups were classified as: short hospital stay (trajectory 1: n=232; 20%); intermediate hospital stay (trajectory 2: n=265; 23%); intermediate hospital stay with discharge limitations (trajectory 3: n=337; 29%); prolonged hospital stay (trajectory 4: n=222; 19%); and death in acute phase (trajectory 5: n=108; 9%).

#### *Vaccination data collection*

Vaccination data was captured in the four convalescent surveys. As enrollment started in March 2021, there was no vaccine data collection built into the acute phase of the study, so data for participants vaccinated prior to the convalescent phase is limited. In the surveys, participants were asked if they had received a vaccine since the last visit, the brand of vaccine, and the date of vaccination.

***Antibody titers***

Anti-SARS-CoV-2 spike, and receptor binding domain (RBD) antibodies were measured by enzyme-linked immunosorbent assay (ELISA) in blood serum as previously described[11]. Readouts were generated in duplicate for serology measures, endpoint titers, and area under the curve (AUC) values. AUC was chosen as the main readout as it considers not only the endpoint titer of ELISA curves but also the magnitude of the signal at each dilution.

***Viral load***

SARS-CoV-2 viral load was assessed by RT-PCR of the viral N1, and N2 genes from nasal swab samples in a centralized laboratory as previously described[1]. Gene PCR cycle threshold (Ct) values were chosen as the main readout.

### MOFA model construction

Multi-Omics Factor analysis is a computational method to identify the principal sources of variation in multi-omics datasets. It employs a variational Bayesian framework, modeling the high-dimensional multi-omics assays as a product of a lower dimensional factor loadings and scores with error. The MOFA factor construction procedure is unsupervised, without knowledge of the outcome variable. In this work, we compared the predictive performance of unsupervised MOFA factors and supervised SPEAR factors. As the SPEAR model construction procedure does not handle missing values, we also employed the MOFA model for missing value imputation prior to constructing the SPEAR model.

We trained separate MOFA models with the MOFA2[12] R package for the Train and Test cohorts, to avoid data leakage between these sets. Both models were obtained initially specifying 250 factors but dropping factors that explain less than 1% of variance in all assays. After this selection the MOFA model on the Train data comprised 156 factors, and the MOFA model on the Test data comprised 48 factors. The following summed explained variance per assay across all factors was obtained: Train – SO: 56.4%, PPT: 19.2%, PPG:45.2%, PMG:59.2%, PGX: 79.2%, BCT: 70.3%. Test – SO: 39.4%, PPT:12.9%, PPG: 44.9%, PMG: 44.2%, PGX: 74.3%, BCT: 63.6%.

### SPEAR model training

***Data imputation***

Signature-based multiPle-omics intEgration via lAtent factoRs[13] (SPEAR) performs supervised dimensionality reduction to identify predictive low-dimensional factors from high-dimensional multi-omics data. SPEAR requires full-rank matrices for constructing a model, thus missing data imputation was first performed using the Train MOFA model described in the previous section.

***SPEAR model training for each response variable***

As SPEAR supports multiple categories of response types (e.g. Gaussian, ordinal, multinomial), we trained multiple SPEAR models considering several LC response variables:

- A two-class (binomial) response of MIN (minimal deficits) or LC (Long COVID).
- A continuous (Gaussian) response using the Physical, Cognitive, Mental, Psychosocial Impact, and Dyspnea population-normalized PROMIS scale scores.

All SPEAR models were trained using 10-fold cross-validation on the Train cohort (with balanced classes within the two-class model) to generate 80 factors (identified as a sufficient rank by

SPEAR). Optimal SPEAR weight (w) was defined as the weight with the lowest overall cross-validated error, resulting in w=1 for the binomial two-class model and w=0 for all Gaussian models.

***SPEAR model selection and evaluation***

To evaluate SPEAR model performance on the Train cohort, we trained a separate lasso classifier for each model using a 10-fold cross-validation procedure. We first compared the PROMIS score SPEAR models using the root mean squared error (RMSE) of the predicted PROMIS scores against the true PROMIS scores. We then took the factor scores from the best performing SPEAR model (SPEAR Physical) as well as the two-class SPEAR model (SPEAR LC), training separate binomial lasso classifiers in a 10-fold cross-validation procedure to predict MIN participants from LC participants. Performance was measured via the average Area Under the Receiver Operating Characteristic (AUROC) scores of 100 bootstrapped model training iterations.

***Computing factor scores for the Test cohort and acute phase visits immune profiling***

SPEAR factor scores for preprocessed testing and acute visit measurements samples were calculated using coefficients derived from the pretrained SPEAR Physical model on the Train cohort.

**Statistical analysis and association analysis**

***Factor scores and geometrical mean scores association***

Factor scores or geometrical mean scores were associated with LC (MIN vs LC labels) and PRO groups by fitting a linear mixed effect model (*lmer* function from the lme4 R package[14]) with the factor score to associate as response variable and endpoint (MIN/LC, PRO group, trajectory group) as covariate, adjusting for sex, discretized age quantile and visit number as fixed effects and enrollment site and participant as random effects using the following formula: *Score ~ endpoint + discretized age quantile + sex + visit number + (1|enrollment site/participant ID)*. We tested for the overall significance of the endpoint term with a goodness of fit Chi-Square test. Associations are shown for each participant in the extended test cohort unless otherwise indicated, which includes Test cohort participants with and without PROMIS Physical score values as well as participants in the Train cohort with no available PROMIS Physical score for any measurement, and thus not included in the SPEAR Physical model training (Figure S3). To assess whether the association was significant after accounting for acute trajectory group, we subset the dataset by visit number and performed the same association, including trajectory group and removing visit number as covariates. P-values were then adjusted across all tests performed on the acute or convalescent visits. To evaluate whether the association was maintained for males and females, we performed the same association after subsetting the dataset to males and females and removing the “sex” covariate from the model.

***Factor scores and geometrical mean scores longitudinal association***

Factor scores or geometrical mean scores were longitudinally associated with LC (MIN vs LC labels), PRO groups, and acute trajectory groups by fitting a smoothing splines generalized additive mixed model (gamm) from the *gamm4* R package with the factor score or analyte to associate as response variable and days after admission and endpoint (MIN/LC label, PRO group, or trajectory group) as covariates, adjusting for sex and discretized age as fixed effects and enrollment site and participant ID as random effects using the following formula: *Score ~ endpoint*

*+ discretized age quantile + sex + s(event date, bs = "ts") + s(event date, bs = "ts", by = endpoint) + (1|enrollment site/participant ID)*. We tested for the overall significance of the endpoint term with a goodness of fit Chi-Square test. Associations are shown for each participant and visit in the extended test cohort unless otherwise indicated, which includes Test cohort participants with and without PROMIS Physical score values as well as participants in the Train cohort with no available PROMIS Physical score values for any measurement, and thus not included in the SPEAR Physical model training. To evaluate whether the recovery factor was able to discriminate between MIN vs LC within participants with the same acute trajectory group, we performed the same longitudinal association after subsetting by trajectory group (Figure S4C).

***Blood CyTOF cell frequencies longitudinal association***

Blood CyTOF cell frequencies were associated with the recovery factor by fitting a linear mixed effect model (lme function from the nlme R package) with the factor score as the response variable and individual cell subset frequency as a covariate, adjusting for sex and discretized age as fixed effects and enrollment site and participant ID as random effects using the following formula: *Score ~ cell frequency + sex + discretized age quantile, random = ~1|enrollment site/participant ID*. We tested for the significance of the cell subset frequency with a Wald test. The acute and convalescent cell type associations used the combined train and test splits for those time points. The model coefficient of each cell type was quantile normalized for comparison.

***Effect of vaccination on factor scores***

To assess the effect of vaccination on the factor scores, we fit a linear mixed effect model (*lmer* function from the lme4 R package[14]) with the factor score as the response variable and endpoint (vaccinated yes/no, within 3 weeks after vaccine dose yes/no) as a covariate, adjusting for sex, discretized age quantile and visit number as fixed effects and enrollment site and participant as random effects using the following formula: *Score ~ endpoint + discretized age quantile + sex + visit number + (1|enrollment site/participant ID).* We tested for the overall significance of the endpoint term with a goodness of fit Chi-Square test.

**Factor annotation and enrichment analysis**

Gene Set Enrichment Analysis (GSEA) of the multi-omics factors was performed as previously described[7]. Briefly, the KEGG, Molecular Signatures Database (MSigDB), and Subpathway publicly available knowledgebases were used for functional enrichment.

- KEGG[15]: KEGG pathways and their corresponding gene and metabolite mappings were extracted using the KEGG REST API (Release 102.0), and were used for SO, PMG, PPT, PPG and PGX assay annotation.
- Hallmark[16]: Hallmark pathways were extracted from the MSigDB database (Homo sapiens) using msigdbr (version 7.5.1), and were used for SO, PPT, PPG and PGX assay annotation.
- Subpathway: the subpathway database provided by Metabolon was used for metabolite annotation (PMG assay)[17–19].

GSEA was conducted on each of the multi-omics assays based on the SPEAR Physical model's Factor loadings, which are a measure of the model's ranking of relative importance for each feature for predicting the PROMIS Physical score (response variable). The clusterProfiler[20] R package was used for GSEA computation. We then computed a multi-omics joint p-value from assays which contained non-zero features in each pathway using the Cauchy combination test[21], which is robust to the dependence structure of the underlying p-values to be combined.

#### Geometric mean score

Geometric mean scores were calculated as previously described[22]. First, per-sample scores were calculated from log-transformed analyte expression values by taking the difference between the geometric mean of positive signature analyte values and the geometric mean of negative signature analyte values. The "Hallmark Heme Metabolism" and "Androgenic Steroids" signatures consisted of the leading edge features from the corresponding recovery factor enrichment analysis results. The "Significant SPEAR Analytes" signature was constructed using all multi-omics analytes identified as significantly contributing to the recovery factor, indicated by a SPEAR-assigned posterior probability >= 0.95 (rounded to two significant figures). Finally, the "Combined" signature used the union of the analytes from "Hallmark Heme Metabolism", "Androgenic Steroids", and "Significant SPEAR Analytes" signatures, conserving analyte directionality across signatures.

#### Heme Metabolism validation in the Hanson et al. (2024) cohort

We accessed publicly available filtered and batch corrected gene counts and LC symptom group designations from Hanson et al[23]. The analysis included n=38 participants with persisting symptoms (PS) and n=49 participants with no persisting symptoms (NPS), with timepoints from 0-90 days post symptom onset. Counts were CPM-normalized using the *cpm* function in edgeR[24]. Participant age was discretized into 5 quantiles for consistency across statistical analyses. Geometric means were calculated using the same set of leading edge genes for Hallmark Heme Metabolism as described in the previous section. Association with LC symptom groups was determined by fitting a linear mixed effect model with the geometric mean as response variable and PASC symptom groups as predictor, adjusting for sex, discretized age, and time bin (customized range of dates from first report of COVID-19 symptoms) as fixed effects and participant ID as a random effect using the following formula: *Score ~ endpoint (PS vs. NPS) + discretized age quantile + sex + time bin + (1| participant ID).* Significance of the LC symptom group term was determined with a goodness of fit Chi-Square test.

#### Machine learning clinical models comparison

##### *SPEAR acute and convalescent models*

Lasso classification models to classify participants according to the LC status (LC vs MIN) were constructed using the SPEAR factor scores for each participant at each of the visits (SPEAR Visit 1-10) were constructed on the Train cohort. Additionally, models including the average of the SPEAR factor scores at the acute phase (SPEAR acute) and convalescent phase (SPEAR conv) were compared. Whenever specified, the clinical features from the clinical models described below were included during model training.

##### *SPEAR significant analytes model*

To compare the performance of the SPEAR models utilizing the factor scores models derived from to the full analyte model to the set of SPEAR significant analytes, we constructed a lasso classification model to classify participants according to the LC status utilizing the average values of the 26 SPEAR significant analytes (analytes with SPEAR posterior probability >= 0.95) during the convalescent phase.

#### *Clinical model construction*

A lasso classification model using baseline clinical measurements, denoted as the clinical model, was constructed on the Train Cohort to classify participants according to the LC status for comparison with the SPEAR acute and convalescent models. The clinical models included the following features: age at enrollment, sex, body mass index (BMI), length of hospital stay, Sequential Organ Failure Assessment (SOFA) score, mean values of Spike IgG levels and viral load (N1 Ct) during the acute phase, and the presence of comorbidities including hypertension, diabetes, chronic cardiac disease, chronic kidney disease, malignant neoplasms, chronic neurological disorders, liver disfunction/failure, history of transplants, smoking/vaping, asthma, respiratory diseases other than asthma, substance use, HIV infection, and the total number of comorbidities.

#### *Model training, evaluation, and performance comparison*

All the models mentioned above were evaluated on a 10-fold cross-validation setting on the Train Cohort. Bootstrapping of N=100 models trained updating the random seed for the fold creation was utilized to compare the average area under the ROC curve (AUROC) performance on the cross-validation folds. The average AUROC values across bootstrapped samples were used for plotting and computing the statistical significance of performance differences across models, employing a t-test, and adjusting the p-values when relevant with the Benjamini-Hochberg approach.
